# Amyloid/RNA Condensates with Germ Granule Characteristics Are Present in Rat Epididymal Spermatozoa and Exhibit Prion-Like Propagation Activity

**DOI:** 10.64898/2026.10.30.763233

**Authors:** Kimberly Garza, Uyen Tran, R. Bryan Sutton, Gail A. Cornwall

## Abstract

Environmental exposures to the germline can alter its epigenome resulting in new traits being acquired by offspring. While the epigenetic carrier in mammalian spermatozoa remains unknown, studies in *C.elegans* show that amyloids and sperm-specific populations of germ granules— phase-separated biomolecular condensates enriched in RNA-binding proteins and small noncoding RNAs— transmit heritable epigenetic phenotypes to descendant cells. Here we discovered that rat epididymal spermatozoa contain germ granule-like condensates in the head perforatorium, a cytoplasmic derived structure that participates in early post-fertilization events. The perforatoria condensates contain evolutionarily conserved germ granule markers and small noncoding RNAs and are enriched in amyloid, including those of the clinically relevant amyloidogenic proteins androgen receptor (AR) and TDP43. Further, perforatoria extracts exhibit prion-like propagation behavior by functioning as molecular seeds to template amyloid assembly in recombinant AR and TDP43. We propose that nature has conserved sperm germ granule-like structures throughout evolution to allow an organism to respond and adapt to a changed environment by passing environmentally encoded information to the embryo via a non-Mendelian protein-based mechanism of inheritance.

## INTRODUCTION

Germ cells are uniquely tasked with preserving and transmitting the heritable genome across generations, thereby ensuring species continuity. However, environmental stressors can alter the germline epigenome, affecting offspring health and, in some cases, producing phenotypic changes that are transmitted across generations, including to descendants that were never directly exposed to the original stressor. Unlike genetic alterations, which involve changes in DNA sequence, these heritable effects are thought to arise through epigenetic mechanisms, including alterations in germ cell DNA methylation, chromatin remodeling, and small noncoding RNAs.

Since the first reports of transgenerational epigenetic inheritance in a rat model in 2005 ^1^, an expanding body of work has described heritable phenotypic effects following exposure to diverse environmental challenges during fetal development or adulthood. Endocrine-disrupting chemicals, nutritional perturbations—including caloric restriction, high-fat diets, and folate deficiency—as well as tobacco smoke and alcohol have each been associated with phenotypes emerging in subsequent generations. Reported outcomes span immune, renal, prostate, cardiovascular, metabolic, pulmonary, neural, behavioral, and reproductive functions, with effects observed in both male and female descendants ^2^. Despite accumulating evidence, the mechanisms underlying transgenerational epigenetic inheritance remain incompletely understood. It is still unclear how an environmental perturbation becomes encoded as heritable information in the germline and equally unclear is the identity of the molecular carrier—or combination of carriers—that conveys this information across generations.

Amyloids are proteins that self-assemble into highly ordered cross-β-sheet, and cross-α- structures. While amyloids have been typically associated with neurodegenerative disease and prionopathies, they carry out biological roles in normal cells and tissues in species from yeast to human. These structures are referred to as functional amyloids and have diverse biological roles including as structural scaffolds, storage sites, molecular switches, and antimicrobial agents ^3–8^.

Although the classic definition of amyloid includes cross-β-sheet structures typically assemble into fibrils, modern amyloid research has revealed that cross-β assemblies in biological systems display a variety of structural states including ball-like oligomers, protofibrils, branched matrices, and films. For clarity we use the term amyloid here as structures with cross-β architecture.

We and others have established that functional amyloids perform key roles in the reproductive tract. Functional amyloids are present in the sperm acrosome and the egg zona pellucida as structural scaffolds where they likely carry out roles in early sperm-oocyte interactions ^9–13^, amyloids form an extracellular matrix-like structure around spermatozoa in the epididymal lumen and perform host defense functions to protect the male germline ^8,14^, and amyloids in semen contribute to sperm quality control by promoting the preferential clearance of damaged and apoptotic sperm by macrophages ^15^.

Amyloids also play roles in the inheritance of new traits ^16^. In yeast environmental exposures can induce proteins to switch from their native fold into amyloids (prion switch) resulting in altered protein function that allows the cell to adapt to the changed environment ^17^. An intriguing characteristic of yeast amyloids/prions is their ability to pass on heritable phenotypes, with self- templating prion conformations being propagated through multiple rounds of cell division and transmitted to meiotic progeny ^18^. Thus, yeast prions represent a protein-based mechanism of epigenetic inheritance through which environmental conditions can promote the emergence of novel phenotypes that are maintained across generations.

Recent studies show a similar mechanism of inheritance in the hermaphroditic nematode worm *C.elegans.* Physiological amyloids in the wild-type worm germline were necessary for germ cells to differentiate to both gametic sexes and were stably inherited allowing each generation to first produce spermatozoa before switching to oogenesis ^19^. This study was the first evidence of protein-based inheritance in an animal raising the possibility that a similar mechanism of inheritance might also occur in higher organisms including mammals.

Independent studies in *C. elegans* demonstrated that cytoplasmic germ granules—phase- separated biomolecular condensates enriched in RNA-binding proteins and small noncoding RNAs that regulate gene expression, protect genome integrity, and maintain germ cell identity— serve as carriers of paternal epigenetic inheritance (PEI granules) by transmitting newly acquired phenotypes to subsequent generations ^20^. This report provided the first evidence of distinct germ granule/condensate populations in mature spermatozoa and revealed their vital role in epigenetic inheritance. Although studies were not done to determine if a changed protein structure within the PEI granules was the epigenetic carrier, the heritable phenotype was passed on to descendant cells via WAGO3, a member of the argonaute family which is known to undergo liquid-liquid phase separation and to form amyloid-like structures ^21^. Indeed, the heritable transmission of the phenotype also required a BTB domain protein responsible for maintaining the structural architecture of *c. elegans* germ granules, demonstrating that the integrity of these ordered protein assemblies is essential for germ granule function and the inheritance of epigenetic phenotypes. Although germ granules are in the germline of all animal species and play essential roles during spermatogenesis, they are thought to be absent from mammalian spermatozoa.

Our previous studies of functional amyloids in mouse spermatozoa showed they contributed to a stable scaffold in the acrosome, a structure which is necessary for spermatozoa to penetrate the zona pellucida, an extracellular matrix that surrounds the oocyte ^11^. Because the acrosome undergoes a well-orchestrated exocytotic event known as the acrosome reaction during which acrosomal contents are released to facilitate sperm passage through the zona pellucida, amyloids within the acrosome would not likely end up in the fertilized oocyte to act as epigenetic carriers. Therefore, we carried out studies here to determine if amyloids are present in sperm-derived cytoplasmic structures that are delivered to the oocyte during fertilization, providing a mechanism for protein-based inheritance in mammals.

We discovered amyloid condensates in fertilization-competent rat cauda epididymal spermatozoa that exhibit the molecular characteristics of germ granules. These condensates are localized to the sperm perforatorium, a juxtanuclear, cytoplasm-derived structure that is delivered into the oocyte at fertilization and participates in post-fertilization and early zygotic events. The perforatorial condensates contain evolutionarily conserved germ granule components, including the germline determinant DDX4/VASA, the piRNA pathway proteins PIWIL1 and PIWIL4, and the scaffolding proteins TDRD6 and TDRD7, which are required for germ granule assembly. They are also enriched in small noncoding RNAs, including piRNAs, tRNA-derived small RNAs (tsRNAs), and microRNAs, suggesting the presence of functional piRNA- and RNA interference- related pathways that contribute to heritable genome protection and epigenetic inheritance. Germ granule proteins are also in the amyloid-rich centrosomal region of the sperm tail, another cytoplasmic structure that is delivered to the oocyte at fertilization. However, the pronounced ubiquitination of this region and the absence of detectable RNA suggest that these assemblies are less likely to be functional and may represent remnants of the chromatoid body, the mammalian germ granule that functions during spermatogenesis but undergoes disassembly during its later stages.^22^ ^23^

The perforatorial amyloid condensates also contain the DNA/RNA-binding proteins androgen receptor (AR) and TDP-43, both of which harbor prion-like domains ^24,25^ and were detected in their amyloid conformations, consistent with biological functions for these assemblies. Importantly, perforatorial extracts exhibited prion-propagating activity, templating assembly by recombinant monomeric AR and TDP-43. These findings suggest that environmentally induced protein conformations—and the biological information they encode—could be transmitted to offspring. While further studies are needed to establish if the germ granule-like condensates and their amyloids transmit heritable phenotypes following an environmental exposure, our results provide a new perspective of the breadth and potential significance of paternal contributions to the embryo including the possible role of a distinct nongenomic, protein-based mechanism of inheritance.

## RESULTS

### Rat cauda epididymal spermatozoa contain amyloid structures in the perforatorium and centrosomal region

We used established reagents that specifically recognize cross-β-sheet architecture to distinguish amyloid structures in rat cauda epididymal spermatozoa. Cauda epididymal spermatozoa are functionally mature having acquired progressive motility and fertilizing ability during transit through the epididymal tubule and are stored in the cauda/tail region of the epididymis until release with mating (Fig 1A). Immunofluorescence analysis using the anti-amyloid antibody A11 (detects immature/oligomeric amyloid) revealed several distinct sites that were labeled including punctate structures in a juxtanuclear region of the rat sperm head known as the perforatorium- a hook structure that extends past the nucleus and is part of the perinuclear theca^26^-(white arrows mark the proximal and distal end of the perforatoria), in distal regions of the perinuclear theca (white arrowheads), a punctate region in the sperm neck/midpiece (yellow arrow), and the sperm midpiece/tail (Fig 1B,C). To more clearly define amyloid localization, intact sperm were separated into heads and tails. Strong anti-amyloid OC (detects mature amyloids including fibrils) and A11 antibody immunoreactivity were detected in the punctate structures in the sperm head perforatorium and perinuclear theca (Fig 1D), while in the isolated tails A11 immunoreactivity was localized along the midpiece and to the centrosomal region at its proximal tip (Fig 1D, white arrow). This region was confirmed as the centrosome using an antibody against speriolin, a sperm-specific centrosomal marker (Fig 1E). ^27^ Although most of the sperm cytoplasm is discarded as residual bodies during release of spermatozoa from the testis and then in cytoplasmic droplets during epididymal transit, the perforatorium/perinuclear theca is a highly insoluble, detergent-resistant remnant of cytoplasm that is intimately associated with the nucleus and that lies in the narrow space between the nucleus and acrosome (Fig 1C). Though the specific functions of the perforatorium remain unclear, it, along with the rest of the perinuclear theca, is delivered into the oocyte during fertilization and is thought to participate in early post- fertilization events, including the release of paternal factors that contribute to oocyte activation ^28–30^. The centrosome is also a highly specialized remnant of cytoplasm that in most species delivers paternal centrioles and associated pericentrosomal material to the embryo for the first mitotic spindle ^31^. Thus, the embryo inherits the architecture of the paternal centrosome. While rat spermatozoa do not contribute a conventional centrosome to the zygote, paternal sperm centrosomal components may nevertheless transmit molecular information to the oocyte. Indeed, mouse sperm speriolin is delivered to the oocyte at fertilization and remains associated with the paternal pronucleus during early zygotic development, providing evidence for its paternal transmission beyond the sperm stage ^27^.

**Figure 1.**
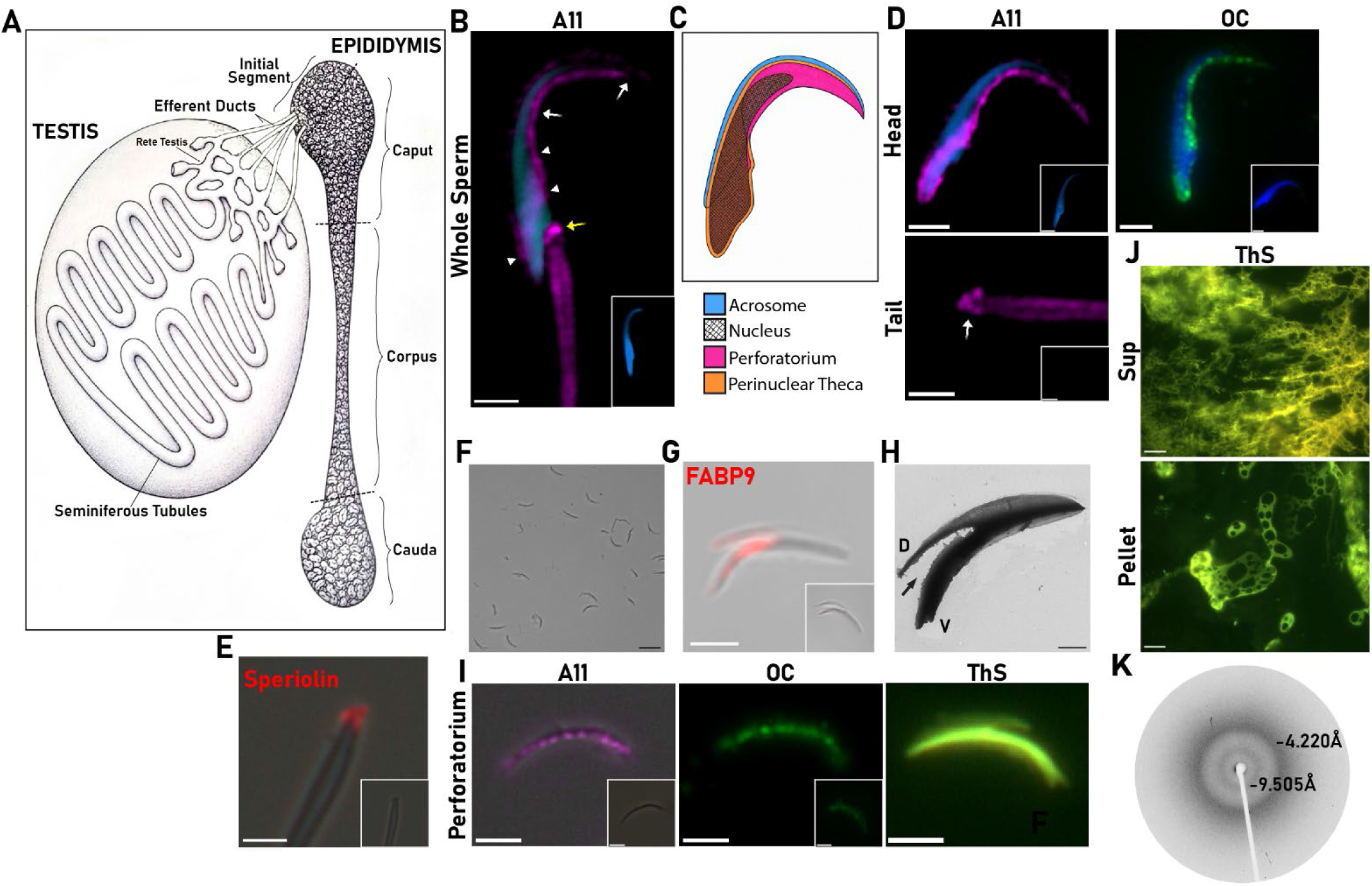
Amyloids are present in the rat cauda epididymal sperm perforatorium and centrosomal region. **A)** Schematic diagram of the rodent testis and epididymis. Adapted from Robaire and Hinton. 1988 ^78^. **B)** Anti-amyloid A11 antibody staining (magenta fluorescence) in intact rat cauda epididymal spermatozoa. White arrows, proximal and distal end of the perforatorium; white arrowheads, perinuclear theca; yellow arrow, sperm neck region. Blue fluorescence, DAPI staining of nucleus. Scale bar, 2.5 μm. Inset, control IgG. Images shown are representative of spermatozoa from n=3 rats. **C)** Schematic diagram of the sperm head regions. **D)** Anti-amyloid A11 (magenta fluorescence) and OC (green fluorescence) antibody staining in isolated rat cauda epididymal sperm heads and tails. Blue fluorescence, DAPI staining of nucleus. White arrow, centrosomal region. Inset, control IgG. Scale bar, 2.5 μm. A11 n=3 rats; OC n=2 rats. **E)** Anti-speriolin (red fluorescence) staining indicating centrosomal region in the sperm tail. Inset, control IgG., Scale bar, 2.5 μm. n=2 rats. **F)** DIC image of isolated perforatoria. Scale bar, 2.5 μm. **G)** Anti-FABP9 (red fluorescence) staining of isolated sperm perforatorium superimposed on DIC image. Inset, control IgG. Scale bar, 2.5 μm. n=4 rats. **H)** Negative stain transmission electron microscopy of an isolated perforatorium showing D, dorsal and V, ventral prongs. Arrow indicates where sperm nucleus would insert. Scale bar, 1 μm. n=4 rats. **I)** Anti-amyloid A11 (magenta fluorescence) and OC (green fluorescence) antibody and thioflavin S (ThS) staining of isolated sperm perforatoria. Inset, control IgG, Scale bar, 2.5 μm. A11 n=5 rats; OC n=1 rat; ThS n=3 rats. **J)** Thioflavin S staining of supernatant (sup) and pellet fraction of an SDS/DTT extract of isolated perforatoria. Scale bar, 40 μm. n=4-5 rats. **K)** X-ray diffraction of isolated rat cauda epididymal sperm perforatoria showing reflections indicative of cross-β structure. n=2 rats.

To more fully examine the amyloid positive puncta, we isolated the perforatoria from rat cauda epididymal sperm heads using a protocol modified from that of Protopapas et al.^28^ This included sonication of purified sperm heads followed by sucrose density gradient centrifugation to isolate the perforatoria from sperm nuclei. The purification scheme resulted in a sample composed of ∼ 90% perforatoria, 8% tail fragments, and 2% nuclei (Fig 1F). Immunostaining with an antibody against FABP9, a perforatorium marker, confirmed that the puncta-containing structure was the perforatorium, showing strong labeling throughout most of this region (Fig 1G). Negative stain TEM of the isolated perforatoria revealed a three-pronged (one dorsal and two ventral prongs) structure as described previously ^28^ with the image in Fig 1H showing a perforatorium on its side. The sperm nucleus inserts into the V-shaped cavity formed by the perforatorium prongs allowing an intimate association and communication between the perforatorium and nucleus (Fig 1H, arrow). Immunofluorescence analysis revealed a line of A11 and OC antibody positive puncta along the ventral prongs of the isolated perforatoria showing that the puncta have amyloid properties (Fig 1I). This was confirmed using the cross-β sheet dye thioflavin S which also indicated the presence of amyloid throughout the perforatorium, although the puncta were difficult to resolve because of the intense fluorescent signal (Fig 1I). Exposure to SDS/DTT which partially disrupted the perforatorium also revealed ThS positive amyloid including in film-like structures from the supernatant and pellet fractions (Fig 1J). Lastly, X-ray diffraction, a gold-standard approach, revealed cross-β-sheet reflections in isolated intact perforatoria, consistent with amyloid (Fig 1K). Together these studies show the perforatorium and its puncta contain cross-β sheets assemblies characteristic of amyloid which is in line with the well-described detergent- resistant properties of the structure^26^. Our detection of amyloid immunoreactivity in the centrosomal region is also in agreement with the centrosome being a biological condensate ^32–34^. We were unable to satisfactorily purify centrosomes from the sperm tail for X-ray diffraction analyses.

### The rat sperm perforatoria puncta and centrosomal region contain germ granule markers

Transmission electron microscopy studies of the ultrastructure of the sperm perinuclear theca and perforatorium from several species including rat have shown that these structures appear as a uniformly electron-dense material with no obvious membranes delimiting specific populations ^26,35^;^36^ ^37^. These data suggest that the amyloid positive puncta in the perforatorium are not vesicles but membraneless organelles, or biological condensates, formed via interactions among proteins with low-complexity regions, intrinsically disordered regions (IDRs), or prion-like domains that drive liquid–liquid phase separation and can mature into gel-like or amyloid states with cross-β architecture. Within the germline these biological condensates are known as germ granules.

To determine if the amyloid puncta in the perforatorium and centrosomal region have the properties of germ granules we first determined if germ granule markers were present. We selected proteins that are conserved across species and that have fundamental germ granule functions including core protein RNA helicase DDX4/Vasa, PIWI pathway proteins PIWIL1/PIWIL4, and scaffold Tudor domain proteins TDRD6 and TDRD7. We also examined Argonaute 2 (AGO2) an miRNA/siRNA effector; Dicer an RNase III that processes precursor RNAs into miRNAs and siRNAs, and BTBD18 and GMCL1, human proteins predicted to be most closely related to those needed to maintain the architecture and heritability of the sperm paternal epigenetic inheritance (PEI) germ granules in *C.elegans* ^20^. All proteins play critical roles in spermatogenesis and except for GMCL1, are essential for male fertility in rodent models and likely also human. DDX4, PIWIL1, Dicer, AGO2, TDRD6 and TDRD7 are also components of the germ granule in the mammalian testis known as the chromatoid body that functions during spermatogenesis ^38–40^. As shown by immunofluorescence analysis BTBD18, GMCL1, Dicer, DDX4, PIWIL1, TDRD6 and TDRD7 localized to the perforatoria condensates both in intact cauda epididymal sperm heads and isolated perforatoria. BTBD18, DDX4, and PIWIL1 were also present in the centrosomal region and, along with Dicer, in small puncta or a line of staining in the region adjacent to the base of the nucleus which may represent the perifossal zone, a specialized, but uncharacterized, structural junction between the sperm head and tail (Fig 2, white arrow). AGO2 was detected in this region as small puncta while PIWIL4 only showed staining in the centrosomal region. The specificity of the antibodies for their antigens was determined by using control IgGs or peptide antigens for blocking the antibody (Fig 2, inset).

**Figure 2.**
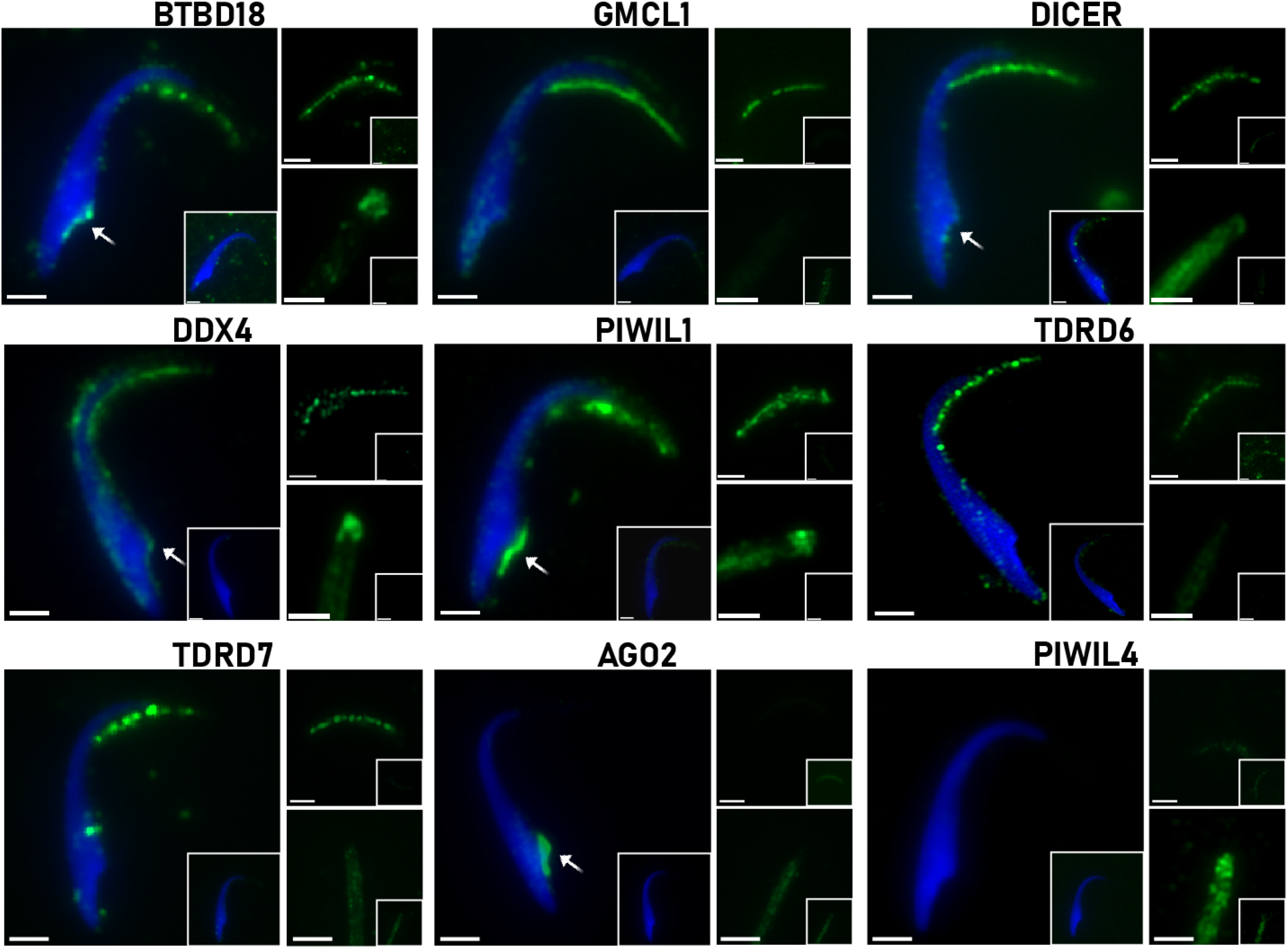
Germ granule markers localize to the rat cauda epididymal sperm perforatoria puncta and centrosomal region. Isolated cauda epididymal sperm heads, purified perforatoria (upper right) and isolated tails (lower right) are shown. Inset, control IgG or peptide blocked Ab. Scale bar, 2.5 μm. White arrows indicate antibody staining in the perifossal zone. n=2-5 rats.

Confocal microscopy and 3D imaging of the perforatoria using Zen software showed that two distinct rows of germ granule marker positive condensates were often present running along the edges of the ventral prongs as shown for DDX4, PIWIL1, TDRD6, BTBD18, and GMCL1 (Fig 3A, Bi-vi). The perforatorium labeled with the anti-Dicer antibody showed only one row of condensates. Indeed, for each germ granule marker and within each cauda epididymal sperm preparation, there was heterogeneity in the distribution of condensates where some sperm contained two positive rows, some only one row and some sperm contained one complete row and a second partial row of condensates. Analysis of the Z-stacks showed that for each protein some were in condensates that aligned within the row as shown for Dicer in Fig 3A where the perforatorium is shown on its side, while in other sperm the condensates were not all in the same plane. The condensates were adjacent to, but outside of, the nucleus and were of varying size ranging from ∼220-380 nm diameter with some spherical in shape and others amorphous. DDX4, PIWIL1, TDRD6, and BTBD18 were also detected in a stretch of smaller puncta ∼ 170-250 nm diameter that ran between the distal part of the perinuclear theca and the perforatorium (Fig 3A arrows). Additional studies are needed to determine if these represent distinct condensate populations and if they merge with those in the perforatorium. We note here that because most of the antibodies used were raised in rabbit we were unable to perform experiments to determine if germ granule markers colocalized within the same perforatoria condensates or if condensates have distinct protein sets. Further studies are needed to determine if the variability in germ granule rows reflects functional differences between individual spermatozoa and the number of condensates present, variability in protein composition of individual condensates, or perforatoria that were damaged.

**Figure 3.**
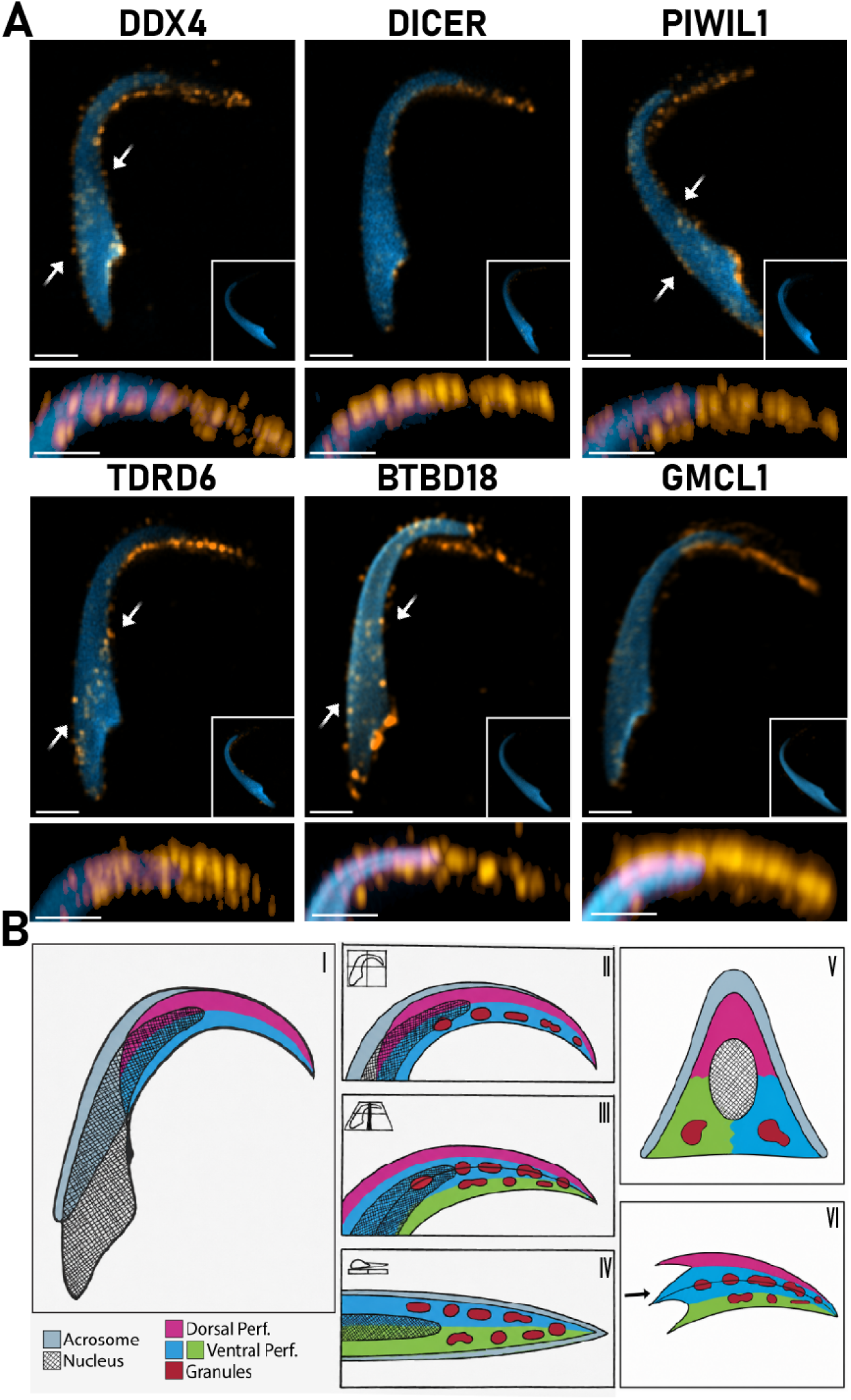
Confocal microscopy and 3D imaging reveal germ granule markers in two distinct rows of condensates. **A)** Immunofluorescence staining (orange fluorescence) of germ granule proteins in isolated cauda epididymal sperm heads. Below, magnified view of perforatoria. Blue fluorescence, DAPI staining of the nucleus. White arrows indicate small puncta that run between the distal part of the perinuclear theca and the perforatorium. Inset, control IgG or peptide blocked antibody. Scale bar, 2.5 μm. n=1-4 rats. **Bi-vi)** Schematic diagram showing the localization of perforatoria condensates in different orientations of the rat cauda epididymal sperm head. Vi arrow, indicates where the nucleus would insert.

Immunoblot confirmed the presence of germ granule markers in spermatozoa and revealed a common protein profile as summarized here and in detail for each protein in SI. A population of mixed germ cells (GC) isolated from the rat testis, containing chromatoid body-positive cells, served as the positive control. When available, blocking peptides were preincubated with the antibody as a control resulting in a marked reduction or elimination of the protein bands and showing the specificity of the antibodies for the germ granule markers (Fig S2). Canonical, full- length proteins (fuchsia arrow) and lower molecular weight isoforms- some previously described and others uncharacterized- (fuchsia asterisks) were detected in testicular germ cells (Fig 4, Fig S1). The isoforms may arise because of several different mechanisms since germ granule proteins are known to exist as alternatively spliced, proteolytically processed, and post- translationally modified forms, mechanisms that may fine-tune their phase separation or aggregation properties as well as their RNA-binding and RNA-processing functions^41–43^. Functionally immature caput (subregions 1-3) and mature cauda (subregions 4-5) epididymal spermatozoa also contained full-length proteins along with multiple lower-molecular-weight isoforms, many of which were more abundant than in testicular germ cells. Examination of isolated cauda epididymal sperm tails, heads and perforatoria showed that most of each germ granule marker localized to the sperm tail and quite often to the insoluble (E3, sarkosyl and E4, urea) fractions consistent with their incorporation into detergent-resistant or aggregated protein assemblies. Further tail populations were often enriched in the lower molecular weight isoforms. In contrast, sperm heads and perforatoria exhibited a protein profile distinct from that of the tails and contained the full-length protein together with lower-molecular-weight isoforms but also higher-molecular-weight immunoreactive species, including material retained in the loading well (fuchsia bracket), indicative of SDS-resistant assemblies. Notably, this distinctive pattern was observed for PIWIL4, TDRD7, BTBD18, and GMCL1 in the isolated perforatoria but was absent from the tail fractions, suggesting that these proteins assemble into unique detergent-resistant complexes within the perforatorium. PIWIL4 and AGO2 were detected in perforatoria by immunoblot using the same antibodies that did not detect protein by immunofluorescence suggesting epitope inaccessibility within the dense assemblies. Together, these findings reveal substantial molecular complexity in germ granule proteins and suggest that their regulation and functions differ between testicular germ cells and mature spermatozoa. Further, our data imply that within spermatozoa the populations of germ granule markers in the tail and perforatoria are biochemically distinct.

**Figure 4.**
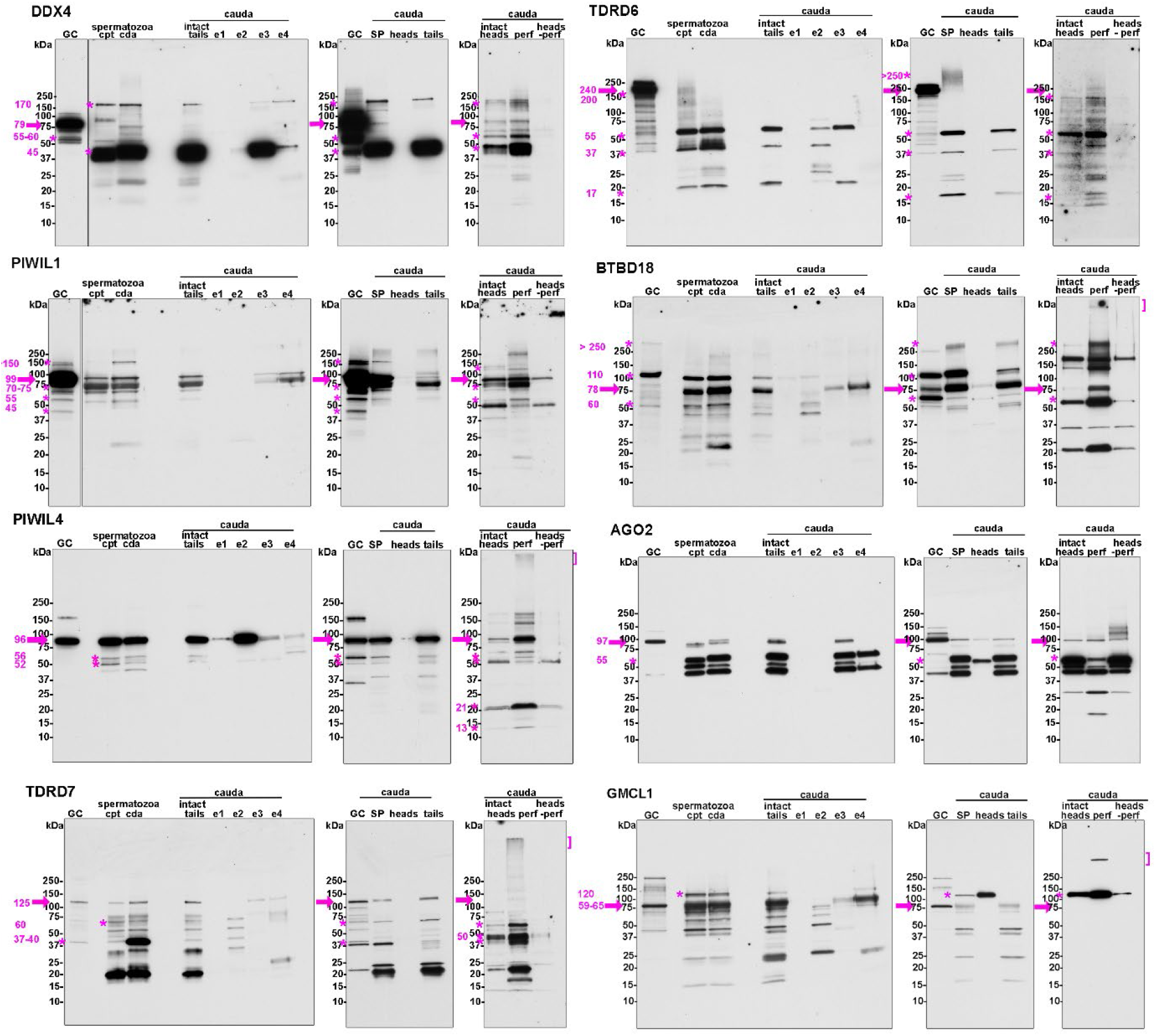
Immunoblot analysis of germ granule markers in epididymal spermatozoa reveals protein profiles distinct from testicular germ cells and differences between sperm heads/perforatoria and tails. Fifty μg testicular germ cells (GC), 1 X 10^6^ intact caput (cpt) and cauda (cda) epididymal spermatozoa, 1.5 X 10^6^ isolated cauda sperm heads, 50 μg isolated sperm tails and 30 μg isolated perforatoria were loaded/lane for each germ granule marker. To examine solubility intact tails (50 μg) were sequentially extracted with E1, Tris buffer; E2, Triton X-100; E3, sarkosyl; E4, 7M urea. Canonical full-length protein, isoforms and larger assemblies are indicated by fuchsia arrow, asterisks, and bracket, respectively. n=2-4 rats. The Dicer immunoblot is shown in Fig S1 and immunoblots with peptide blocked antibodies are in Fig S2. Full details for each protein are in SI text.

### Androgen receptor and TDP43 are present in the perforatoria condensates and centrosomal region

The germ granule proteins examined above contain intrinsically disordered regions (IDRs) which are essential for condensate assembly through multivalent interactions and liquid-liquid phase separation. These dynamic condensates typically exhibit liquid-like material properties. To determine whether proteins containing prion-like domains contribute to condensate formation and thus may promote a more solid, amyloid-like state, we examined androgen receptor (AR) and TDP-43, two clinically relevant amyloidogenic proteins implicated in neurodegenerative disease that have roles in mammalian spermatogenesis.

As shown in intact cauda epididymal spermatozoa, isolated tails, heads, and perforatoria, AR and TDP43 exhibited almost identical patterns of immunostaining with each other and with germ granule markers and were found in the perforatoria condensates (Fig 5A, D, E, F, G, H), centrosomal region (Fig 5B,C), and in the putative perifossal zone adjacent to the base of the nucleus (Fig 5A, F, white arrow). The merged fluorescence signals suggested that AR and TDP43 colocalized within each of these distinct regions in cauda epididymal spermatozoa (Fig 5F). Further, confocal microscopy and Imaris (Oxford Instruments) colocalization analysis revealed the AR and TDP43 colocalized within the perforatoria condensates (Fig 5G,H). A colocalization channel created in the Imaris Coloc module depicting all the voxels that contain both signals showed that 66% of the AR and 56% of the TDP43 signal above the set threshold colocalized. A Pearson’s correlation coefficient of 0.512 suggests a moderate correlation between the two signals, confirming that they are colocalizing in the perforatoria, even if not perfectly (Fig 5G,H).

**Figure 5.**
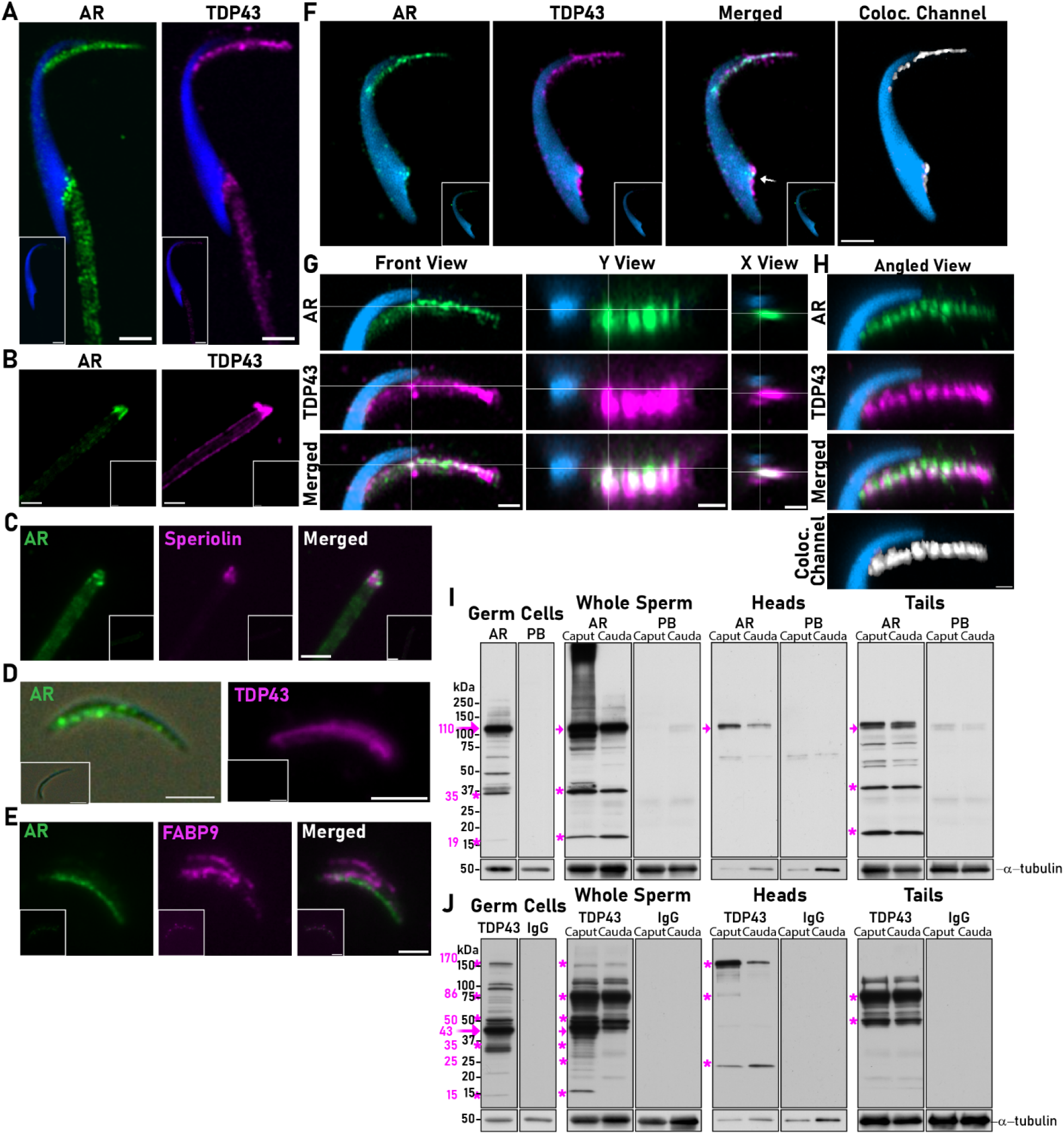
Androgen receptor and TDP43 colocalize in the sperm perforatoria and centrosomal region. **A)** AR (green fluorescence) and TDP43 (magenta fluorescence) are in the perforatorium, centrosomal region and perifossal zone of intact cauda epididymal spermatozoa. Blue fluorescence, DAPI staining of nucleus. Inset, peptide blocked antibody (AR) or control IgG (TDP43). Scale bar, 2.5 μm. AR n=6 rats; TDP43 n=5 rats. **B)** AR (green fluorescence) and TDP43 (magenta fluorescence) staining of the cauda epididymal sperm centrosomal region. Inset, peptide blocked antibody (AR) or control IgG (TDP43). AR n=5 rats; TDP43 n=3 rats. **C)** AR (green fluorescence) colocalizes (merged) with speriolin (magenta fluorescence) indicating AR is present in the centrosomal region. Inset, peptide blocked antibody (AR) or control IgG (speriolin). Scale bar, 2.5 μm. n=2 rats. **D)** AR (green fluorescence) and TDP43 (magenta fluorescence) staining of perforatoria condensates. Inset, peptide blocked antibody (AR) or control IgG (TDP43). Scale bar, 2.5 μm. AR n=5 rats; TDP43 n=7 rats. **E)** AR (green fluorescence) is found in FABP9 (magenta fluorescence) positive structures (merged) indicating AR is present in the perforatoria. Inset, peptide blocked antibody (AR) or control IgG (FABP9). Scale bar, 2.5 μm. n=3 rats. **F)** AR (green fluorescence) and TDP43 (magenta fluorescence) colocalize (merged, Colocalization channel) in the perforatoria condensates and perifossal zone in isolated cauda epididymal sperm heads. Inset, peptide blocked antibody (AR) or control IgG (TDP43). Scale bar, 2.5 μm. **G)** Front, Y-view, X-view, and **H)** angled view showing colocalization (white) of AR (green fluorescence) and TDP43 (magenta fluorescence) in the cauda epididymal sperm perforatoria condensates. Scale bar, 1 μm. n=3 rats. **I) J)** Immunoblot showing full-length (fuchsia arrow) and isoforms (fuchsia asterisks) of **I)** AR and **J)** TDP43 in testicular germ cells (GC), intact caput and cauda epididymal spermatozoa and isolated caput and cauda sperm heads and tails. Fifty μg GC, 1 X 10^6^ intact caput and cauda sperm, 1.5 x 10^6^ isolated sperm heads and 50 μg tails were loaded in each lane. PB, peptide block. IgG, control antibody. Blots were stripped and reprobed with α-tubulin as a loading control. n=4-5 rats.

Immunoblot confirmed the full-length AR (∼110 kDa) was the predominant protein in testicular germ cells, intact spermatozoa, and isolated sperm heads and tails (Fig 5I, fuchsia arrow). In addition to full-length AR, smaller forms including those at 35 kDa and 19 kDa, that appeared to be primarily in the sperm tail, were detected and may represent proteolytically processed forms of the protein (Fig 5I, fuchsia asterisks). The 35 kDa AR protein is thought to be a protease resistant fragment that is present in AR protein in a ligand bound, conformationally altered state, formed during its activation^44,45^. While full-length (43 kDa) TDP43 was the predominant form in testicular germ cells, higher molecular weight TDP43 proteins of ∼86 and ∼170-180 kDa, likely dimers and tetramers, respectively, were the predominant TDP43 proteins in sperm heads and tails (Fig 5J)^46^. The 50 kDa TDP43 may represent phosphorylated TDP43. Several C-terminal fragments that are often found in self-assembled TDP43 oligomers including proteins at 35 kDa, 25 kDa, and 15 kDa were also detected.

### The sperm tail is enriched in mono and polyubiquitinated proteins

The presence of germ granule markers in two distinct amyloid-rich sperm compartments, along with their distinct protein profiles, suggests functional specialization between these populations. Cells possess an aggresomal pathway in which misfolded proteins and amyloid-like aggregates, many of which are ubiquitinated, are transported to aggresomes for subsequent clearance through proteasomal degradation or autophagy. Notably, aggresomes are typically localized to the pericentrosomal region, suggesting that germ granule components associated with the sperm tail centrosomal region may undergo differential processing or turnover compared with those localized to the perforatorium^47^. To investigate whether ubiquitinated proteins are differentially distributed between these sperm compartments, we performed immunoblot analysis using an antibody that recognizes both mono- and polyubiquitinated proteins. Sperm tail fractions, which contain the centrosome, exhibited substantially higher levels of ubiquitinated protein particularly in the insoluble e3 and e4 extracts, compared with isolated sperm head fractions containing the perforatorium (Fig. 6A). These observations were supported by immunofluorescence studies that showed an enrichment of ubiquitinated proteins in the sperm centrosomal region (Fig 6B). Ubiquitin staining only became apparent after exposure of the sperm tail fractions to trypsin suggesting the ubiquitin tagged proteins are in complex insoluble assemblies.^48^ Although the ubiquitination status of individual germ granule markers was not determined, these findings further indicate biochemical differences between germ granule-associated assemblies in the perforatorium and centrosomal/tail regions, with those in the latter possibly targeted for turnover through ubiquitin-dependent pathways.

**Figure 6.**
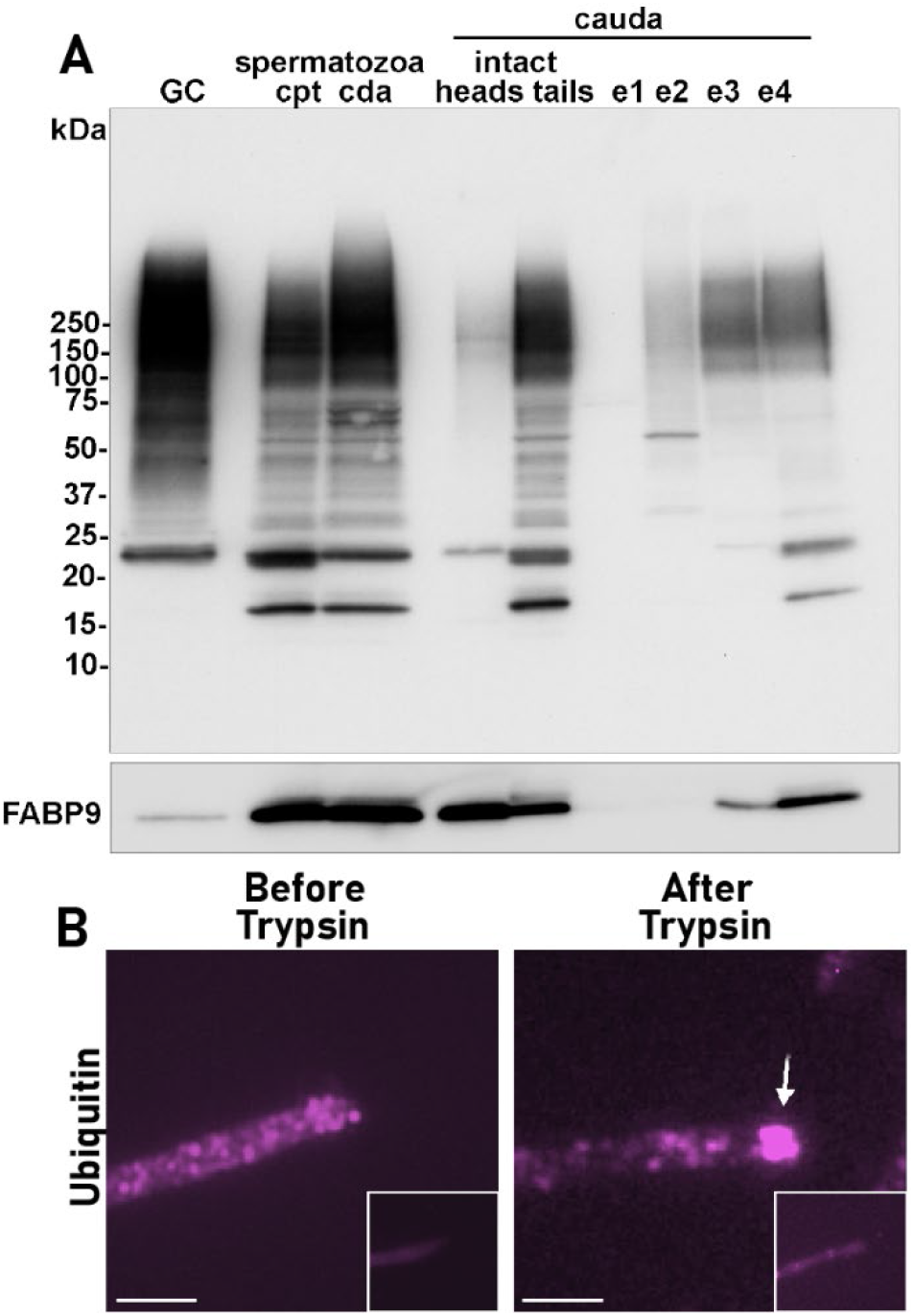
Mono- and poly-ubiquitinated proteins are enriched in cauda epididymal sperm tails/centrosomal region. **A)** Immunoblot of testicular germ cells (GC, 50 μg), 1 X 10^6^ intact caput and cauda epididymal spermatozoa, 1.5 X 10^6^ cauda sperm heads, 50 μg cauda sperm tails and equal proportions of sperm tails extracted in E1, Tris; E2, Triton X-100, E3, sarkosyl, and E4 urea using an anti- ubiquitin antibody. The blot was stripped and reprobed with FABP9, a marker for the peforatorium to show loading of sperm heads. Some FABP9 has been previously reported to also be in the sperm tail as indicated here. ^79^ n=2 rats. **B)** Immunofluorescence analysis showing ubiquitinated proteins are detected in the sperm centrosomal region (white arrow) after exposure to trypsin. Inset, control IgG. Scale bar, 2.5 μm. n=2 rats.

### The perforatoria condensates contain RNA

Germ granules serve as critical hubs for RNA-binding proteins that carry out genome-protective and gene-regulatory functions through interactions with guide RNAs, including piRNAs, miRNAs, and tsRNAs, which direct these proteins to their target transcripts. Accordingly, the presence of RNA is a defining feature of germ granules and is essential for their roles in RNA-mediated regulation^49^. Using RNASelect, an RNA-selective fluorescent nucleic-acid dye, we detected fluorescence within the perforatorium condensates but not in the centrosomal region, consistent with enrichment of RNA within the perforatoria (Fig 7A). Sperm tails that were first exposed to trypsin to help reveal centrosomal components prior to RNASelect staining also did not show fluorescence further suggesting that RNA was not present. Because negative RNA staining in the centrosomal region may reflect limited dye accessibility in this ubiquitin-rich, insoluble compartment, we cannot exclude the presence of inaccessible or low-abundance RNA. However, based on these findings we focused our efforts on characterizing RNA populations in the perforatorium. Although attempts were made to isolate the condensates, this proved difficult due to the insoluble nature of the entire perforatorium. Therefore, we isolated total RNA from pooled populations of intact perforatoria to obtain an overall view of the populations that are present. Separation of the RNA on an SDS-urea gel revealed populations of small RNAs in addition to larger forms, likely mRNA (Fig 7B). SncRNA-seq revealed distinct populations in the perforatoria with the predominant populations being tRNA/tRNA derived fragments (tsRNAs), piRNAs, and circular RNAs-covalently closed RNA molecules (Fig 7C). Surprisingly, miRNAs represented only a small percentage of the total sncRNAs and were mostly present as pre-miRNAs rather than mature miRNAs. Our inability to detect mature miRNAs may be due to their very low representation or because pre-miRNAs may be stored in the perforatoria in an immature state until processing and function in the zygote. Our data are consistent with other reports showing that miRNAs are more abundant in sperm tails while tRNAs are more abundant in sperm heads ^50^. Other populations of sncRNA in the perforatorium were small nucleolar RNAs (snoRNAs) that direct post-transcriptional modification and processing of rRNA, although some can function like miRNAs, and small nuclear RNAs (snRNAs) that are core component of the spliceosome. While piRNAs, tRNA/tsRNAs, and miRNAs have roles in epigenetic inheritance, the contributions of circular RNAs have not been determined. However, based on their unusual stability, several functions have been proposed including roles as protein-binding scaffolds/organization centers for RNA binding proteins/amyloids or as direct regulators of other sncRNAs^51^. Collectively, our experiments suggest that the perforatoria condensates are functional germ granules as they possess the critical regulatory proteins and sncRNA guides to regulate gene expression and protect the genome. In-depth analyses of the sncRNA populations in the perforatoria are needed to identify their potential mRNA targets and functions.

**Figure 7.**
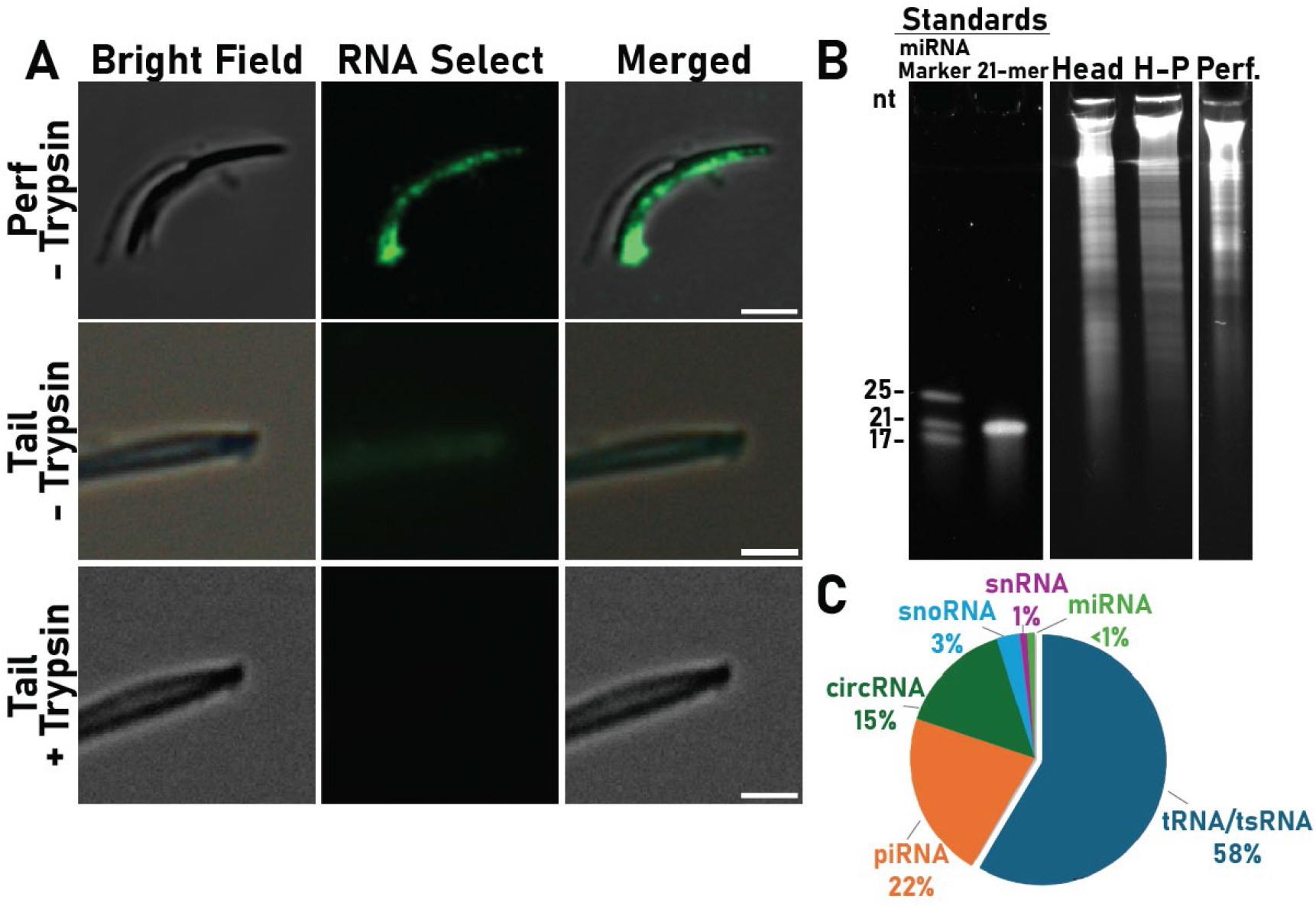
Cauda sperm perforatoria condensates contain RNA. **A)** SytoRNA select staining of isolated perforatoria and tails revealed RNA in the perforatoria condensates but not in the centrosomal region including after prior exposure to trypsin. Fluorescence is superimposed on DIC images. Scale bar, 2.5 μm. n=4 rats. **B)** SDS-urea gel showing RNA distribution in intact cauda sperm heads, heads without perforatoria (H-P) and isolated perforatoria (perf). Intact heads n=4 rats, heads without perforatoria and isolated perforatora, n=1 RNA isolation from perforatoria pooled from 5-6 rats. **C)** Distribution of sncRNA populations in isolated cauda sperm perforatoria. n=3 RNA isolations each generated from perforatoria pooled from 5-6 rats.

### Formation and maturation of perforatoria condensates in spermatozoa

The mammalian chromatoid body disappears from late-stage elongating spermatids in the testis ^23,52^. The fate of chromatoid body proteins and RNAs is unknown but is thought to include dispersion into smaller granules or release into the cytoplasm and elimination by residual bodies. We used PIWIL1, BTBD18, and DDX4 as markers of cauda sperm perforatoria condensates to determine whether condensates are also present in the perforatoria of testicular and immature epididymal spermatozoa from the initial segment (subregion 1), the beginning of the epididymal tubule (Fig 8A). Previous studies in the rat show the perforatorium becomes a distinct structural entity only at the very end of spermiogenesis abruptly acquiring its electron dense properties in late-stage (step 19) elongating spermatids^53^. Because testicular spermatozoa were recovered in low numbers and are more fragile than cauda spermatozoa, isolation of perforatoria was not feasible. Therefore, whole spermatozoa were examined in this experiment. Immunofluorescence analysis and confocal microscopy showed that PIWIL1, BTBD18, and DDX4 were present as condensates in testicular sperm perforatoria (Fig. 8B, white arrows). PIWIL1 was consistently detected in a full row of condensates in the perforatoria while BTBD18 and DDX4 were more variable and sometimes present in incomplete rows. FABP9 was also detected throughout this region, but not as condensates, confirming it as perforatorium (Fig. 8C). Some testicular spermatozoa did not exhibit detectable BTBD18 or DDX4 in the perforatoria. By the initial segment, however, spermatozoa revealed all three proteins in distinct perforatoria condensates, usually as complete rows that became slightly more defined during sperm migration to the cauda, suggesting that additional maturation/consolidation occurs during epididymal transit. A diffuse distribution of PIWIL1, BTBD18, and DDX4 was also detected along the back ridge of the testicular, initial segment, and cauda sperm head (white arrowheads), which may represent protein in other parts of the perinuclear theca or the acrosome. Depending on the protein, the back ridge population varied in intensity between the sperm samples from the three different regions which in initial segment and cauda epididymal spermatozoa may represent remodeling of the sperm surface including the acquisition of proteins from the efferent duct and epididymis. It is possible that in testicular spermatozoa the back ridge population serves as a source of germ granule markers for perforatoria condensate assembly. Compared to intact cauda spermatozoa, isolated cauda sperm heads exhibited little to no fluorescence for PIWIL1, BTBD18, or DDX4 along the back ridge of the sperm head (Fig 2). This reduction in signal may reflect loss of these components during head–tail separation. PIWIL1, BTBD18 and DDX4 were also detected in the sperm neck and in the midpiece but due to background in testicular sperm tails and the challenge distinguishing tail structures from head in intact spermatozoa we were unable to definitively conclude that germ granule markers were in the centrosomal region of testicular and initial segment spermatozoa. Together, our data reveal that the formation of perforatoria condensates occurs in the testis with possibly additional consolidation in post-testicular regions including the efferent ducts that connect the testis to the epididymis. These results imply that exposure of germ granule components to environmental insults during early germline development or in adulthood could induce persistent changes that are carried forward into fertilizing cauda epididymal spermatozoa and to the zygote.

**Figure 8.**
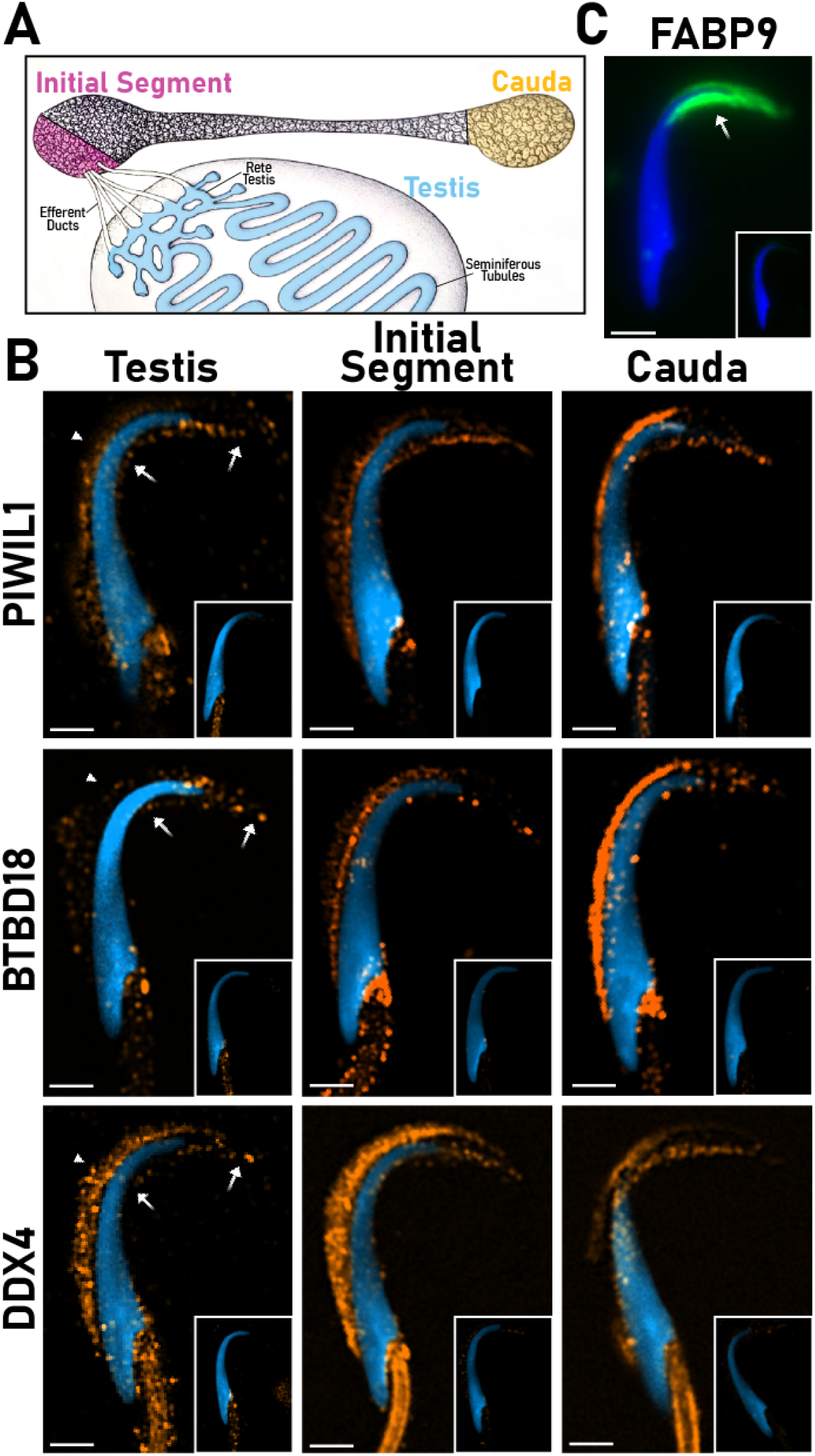
Germ granule-like condensates are in testicular sperm perforatoria. **A)** Schematic diagram of the rodent testis and epididymis showing the regions from which spermatozoa were examined for germ granule markers. **B)** Immunofluorescence analysis showing PIWIL1, BTBD18 and DDX4 in perforatoria condensates in spermatozoa from the testis, initial segment, and cauda epididymis. Inset, peptide blocked antibody. Scale bar, 2.5 μm. white arrows indicate proximal and distal ends of the perforatorium. White arrowheads indicate back ridge staining. n=3-4 rats. **C)** Immunofluorescence analysis showing FABP9 in the perforatorium (white arrow) of testicular spermatozoa. Inset, control IgG. Scale bar, 2.5 μm. n=2 rats.

### AR and TDP43 amyloids are present in rat sperm perforatorium and centrosomal region

Because AR and TDP-43 are amyloidogenic, their presence within the perforatoria condensates and centrosomal region suggests they contribute to the amyloid properties of these structures. Therefore, we used multiple strategies to determine if AR and TDP43 amyloids are present in the perforatorium and centrosomal region. Colocalization studies and confocal imaging showed that AR was present in A11 amyloid positive condensates in the perforatoria (Fig 9A-C).

**Figure 9.**
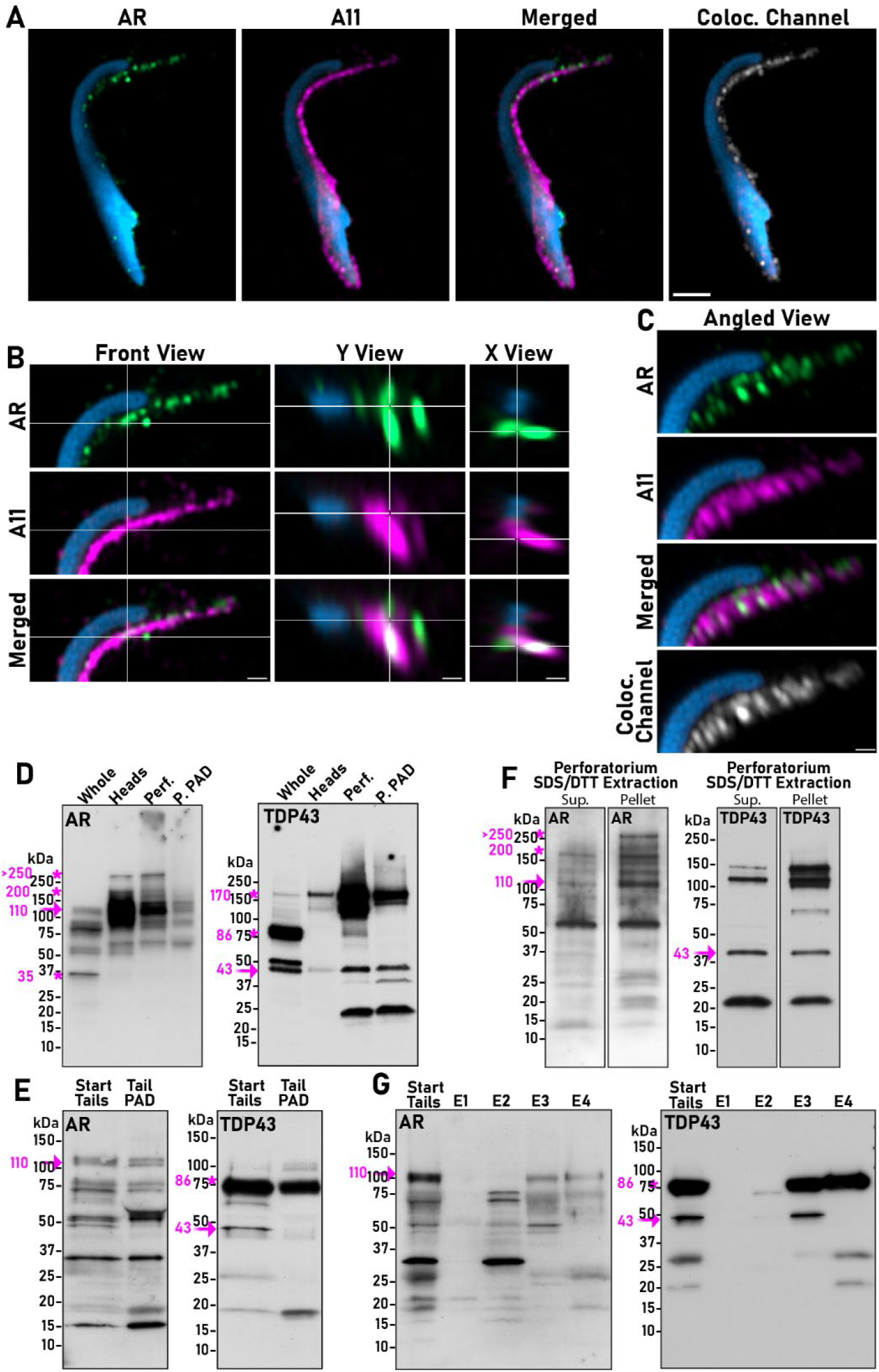
Cauda epididymal sperm perforatoria contain AR and TDP43 amyloids. **A)** Colocalization of AR (green fluorescence) and A11 (magenta fluorescence) in cauda epididymal sperm head perforatorium. Analysis with Imaris Coloc, which generated a new channel where every voxel that contains signal above the set threshold from both channels appears as white, demonstrates that AR and A11 colocalize throughout the perforatorium even if the one channel is at a lower intensity than the other. **B)** Orthogonal views of the perforatorium show a granule of strong colocalization between AR and A11, seen as a white puncta since both channels are present at similar and high intensities. **C)** A perforatorium zoomed-in view angled in the z-plane about 45 degrees shows the 3- dimensional nature of the signal, with the Imaris-generated colocalization channel showing that although not visually obvious (since the intensities of the two channels are not equal/balanced) there are plenty of voxels where both channels colocalize. A-C) n=3 rats. **D)** Immunoblot showing AR and TDP43 in the sperm perforatoria binds to the PAD ligand suggesting amyloid properties. Full-length AR or TDP43 (fuchsia arrow), AR or TDP43 isoforms (fuchsia asterisks). Whole, 1 x 10^6^ intact cauda epididymal spermatozoa; heads, 1.5 x 10^6^ isolated cauda epididymal sperm heads; perf, 35 μg intact perforatorum for AR and 29 μg intact perforatorium for TDP43; perf PAD, 48 μg isolated perforatoria were used for PAD pulldown of AR and 129 μg for TDP43. AR, n=3 rats; TDP43, n=4 rats. **E)** Immunoblot of AR and TDP43 showing protein in the sperm tails bind to the PAD ligand suggesting amyloid properties. Full-length AR or TDP43 (fuchsia arrow), AR isoforms or TDP43 dimers/tetramers (fuchsia asterisks). Fifty μg start tails were loaded and 400 μg were used for PAD pulldown. n=3 rats. **F)** Immunoblot showing AR and TDP43 in the supernatant and pellet fractions of SDS/DTT extracts of perforatoria isolated from cauda epididymal spermatozoa. Full-length AR or TDP43 (fuchsia arrow), AR isoforms or TDP43 dimers (86 kDa), tetramers (170 kDa) ((fuchsia asterisks). Twenty μl each of supernatant and pellet were loaded/lane. AR, n=3 rats; TDP43, n=2 rats. **G)** Immunoblot of AR and TDP43 in cauda epididymal sperm tails (start) and following sequential extraction of tails in E1-E4 buffer. Thirty μl of each extraction and 50 μg of starting nonextracted tails were loaded/lane. AR, n=2 rats; TDP43, n=2 rats.

We also exposed isolated perforatoria and tails to DMSO to help unwind the amyloid, followed by incubation with the PAD (protein aggregation disease) ligand which captures amyloid but not protein monomers. These experiments showed that the intact perforatorium contained full-length AR (∼110 kDa), high molecular weight SDS-resistant forms (∼200 kDa and >250 kDa), and smaller processed forms. After exposure to DMSO and PAD binding, the full-length and processed forms were detected suggesting a population of AR is present in the perforatorium as an amyloid structure (Fig 9D). Similarly, full-length and high molecular weight SDS-resistant forms of TDP43, including a population retained in the well, were the predominant proteins present in the perforatorium and bound to the PAD beads suggesting amyloid conformations (Fig 9D). Incubation of DMSO-exposed cauda sperm tails with the PAD ligand also revealed populations of AR and TDP43 amyloids (Fig 9E).

Since amyloids are often SDS-resistant, we examined the supernatant and pellet fractions after perforatoria were extracted with SDS/DTT. While AR and TDP43 were found in the soluble fraction (supernatant), a proportion of both proteins also remained in the insoluble pellet, including high molecular weight SDS-resistant forms of AR, characteristic of aggregated protein/amyloid (Fig 9F). Correspondingly, the sequential extraction of sperm tails with buffers containing different detergents (E1-E4) showed that most full-length and higher molecular weight forms of AR and TDP43 required sarkosyl (E3) and 7M urea (E4) for extraction, indicative of insoluble populations (Fig 9G).

Because amyloids are also protease-resistant, head and tail extracts were exposed to proteinase K (PK) and the digestion of AR and TDP43 monitored over time. In contrast to recombinant AR and TDP43 monomers which were sensitive to PK and completed digested with 1-2 hrs, AR and TDP43 in the cell extracts were resistant, suggesting their presence in a stable structure such as amyloid. AR proteins in the sperm tail were profoundly more resistant to PK (Fig 10B) than populations in the head (Fig 10A) even in the presence of more than 2x the amount of PK than that used to digest sperm heads. A reprobing of the blots with anti-TDP43 antibody revealed that while TDP43 in cell extracts was also resistant to PK treatment compared to its monomer, it was more sensitive to digestion than AR (Fig 10C, D). Further, the different isoforms of TDP43 exhibited different sensitivities to PK. In the sperm head the 43 kDa form of TDP43 (arrow) was more resistant than the 150 kDa, a functional tetramer that normally exists in cells ^54,55^. The 25 kDa TDP43, a known C-terminal fragment that is highly aggregation prone remained resistant to PK throughout the 4 hr time course. Like AR, all forms of TDP43 in the sperm tail were resistant to PK suggesting highly insoluble structures. Collectively, our data suggest that while perforatoria condensates and the centrosomal/tail region contain AR and TDP43 amyloids, their structural properties differ with those in the centrosomal /tail region more resistant to solubilization than those in the perforatorium.

**Figure 10.**
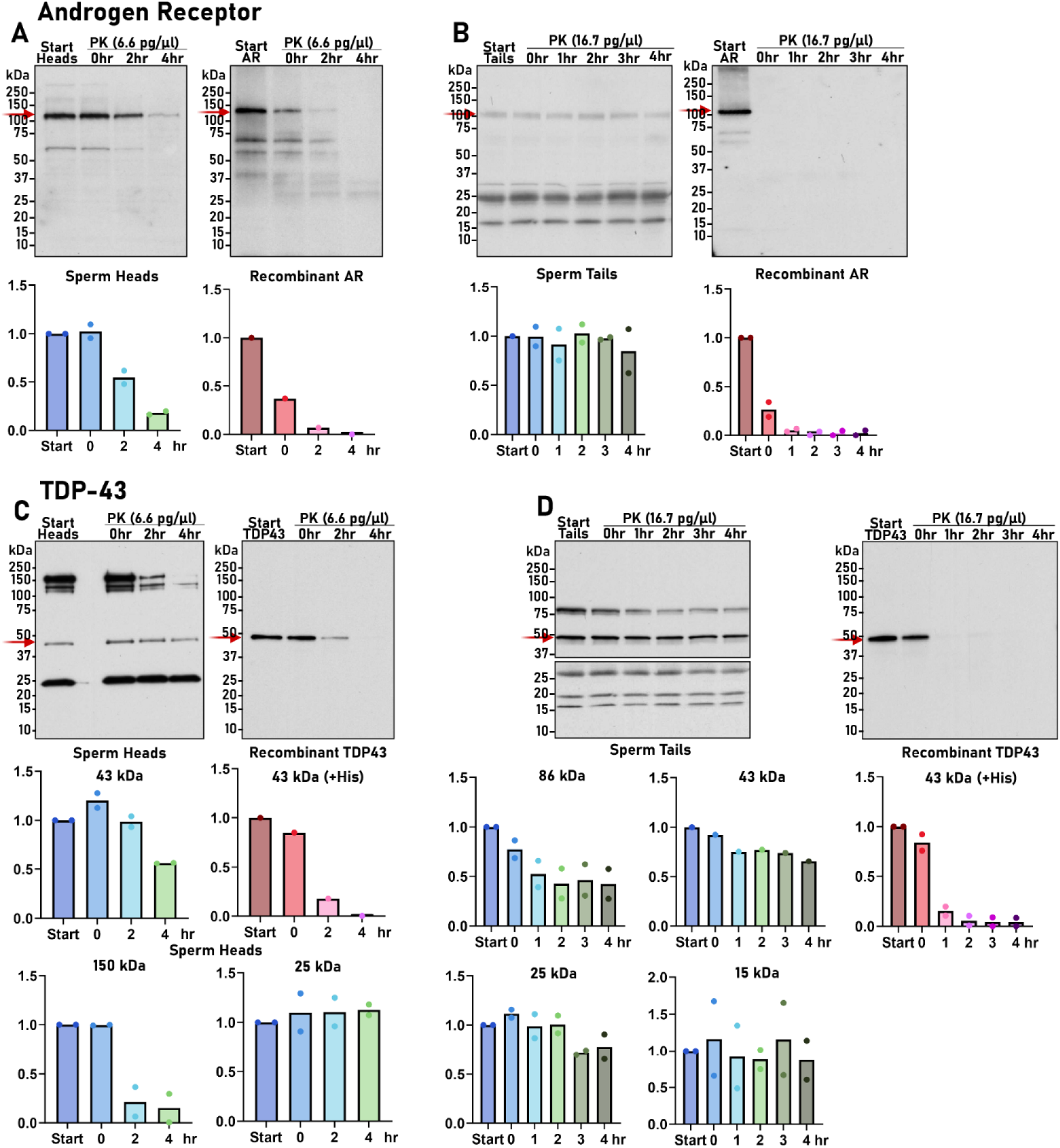
AR and TDP43 in sperm heads and tails are proteinase K resistant. **A) B)** Immunoblot comparing the proteinase K (PK) resistance of AR in **A)** cauda epididymal sperm heads and **B)** cauda epididymal sperm tails with that of recombinant full-length rat AR produced in bacteria in-house. Band intensities for full-length AR (fuchsia arrow) were quantified by Image J and normalized to starting sample (start) that was not exposed to PK and are shown in bar graphs below the blots. The individual points for n=2 experiments (1 rat/experiment) are shown. **C)D)** Immunoblot comparing the PK resistance of TDP43 in **C)** cauda epididymal sperm heads and **D)** cauda epididymal sperm tails with that of recombinant TDP43. Band intensities for full- length (43 kDa, fuchsia arrow), 150 kDa (tetramer), 25 kDa and 15 kDa isoforms were quantified by Image J and normalized to starting sample (start) that was not exposed to PK and are shown in bar graphs below the blots. The individual points for n=2 experiments 1 rat/experiment) are shown. The blot in **D)** showing the lower molecular weight forms of TDP43 in the sperm tail was exposed longer to better visualize the bands. The 15 kDa TDP43 isoform in the sperm tail showed a similar trend in response to PK but was present in different levels between the two experiments. The top row of points represents one experiment while the bottom points indicate the second experiment.

### Perforatoria extracts template amyloid assembly in AR and TDP43

To function as epigenetic carriers, sperm amyloids need to be able to propagate their structures in naïve/monomeric protein allowing transmission of structurally encoded information and potentially new phenotypes. Since we could not isolate condensates from the perforatorium, we used the supernatant of SDS/DTT extracts from normal SD rat cauda sperm peforatoria to determine if they contained molecular seeds/templates to initiate amyloidogenesis in AR and TDP43 monomers. Because of nonspecific interactions between SDS and thioflavin T (ThT) we were unable to observe classic amyloid assembly kinetics using standard ThT assays^56^. However, when the reactions were examined by negative stain transmission electron microscopy (TEM), very different morphologies were observed in the seeded reactions compared to monomer or perforatoria seeds alone. AR or TDP43 monomeric protein alone (in the presence of SDS/DTT) contained little to no structure while the perforatoria seeds (SDS/DTT extracts) had faint granular patches, small, scattered clumps, and sometimes stacked shards of perforatoria material (Fig 11 A, B). In contrast, incubation of AR (AR + seeds) or TDP43 (TDP43 + seeds) with SDS/DTT perforatoria supernatant (seed) resulted in assemblies of branched matrix that were stained with uranyl acetate (Fig 11A, B). Upon higher magnification some of these darkly stained matrices revealed oligomers and protofibrils (Fig 11B, Fig S3B,C). Similar results were observed using several different recombinant AR protein preparations including a full-length rat AR produced in bacteria in-house (Fig 11A,C, Fig S3D), commercially prepared full-length rat AR produced in bacteria (Creative Biomart, Shirley, NY) (Fig S3A,D), full-length AR produced by HEK cells in- house (Fig S3B,D), and N-terminal rat AR lacking the ligand binding domain and produced in bacteria in-house (Fig S3C, D). In addition to branched matrix, films and fibrils were also observed when the commercially prepared AR was seeded with the perf SDS/DTT seed (Fig S3A). For TDP43, a commercially prepared full-length human TDP43 protein was used which exhibits ∼ 96% sequence similarity with rat TDP43 (Fig 11B,D, Fig S3D). In general, TDP43 seeded reactions formed smaller assemblies compared to that with AR, consisting of patches of oligomers/protofibrils and matrices and films that were less heavily stained with uranyl acetate compared to AR seeded reactions (Fig 11B).

**Figure 11.**
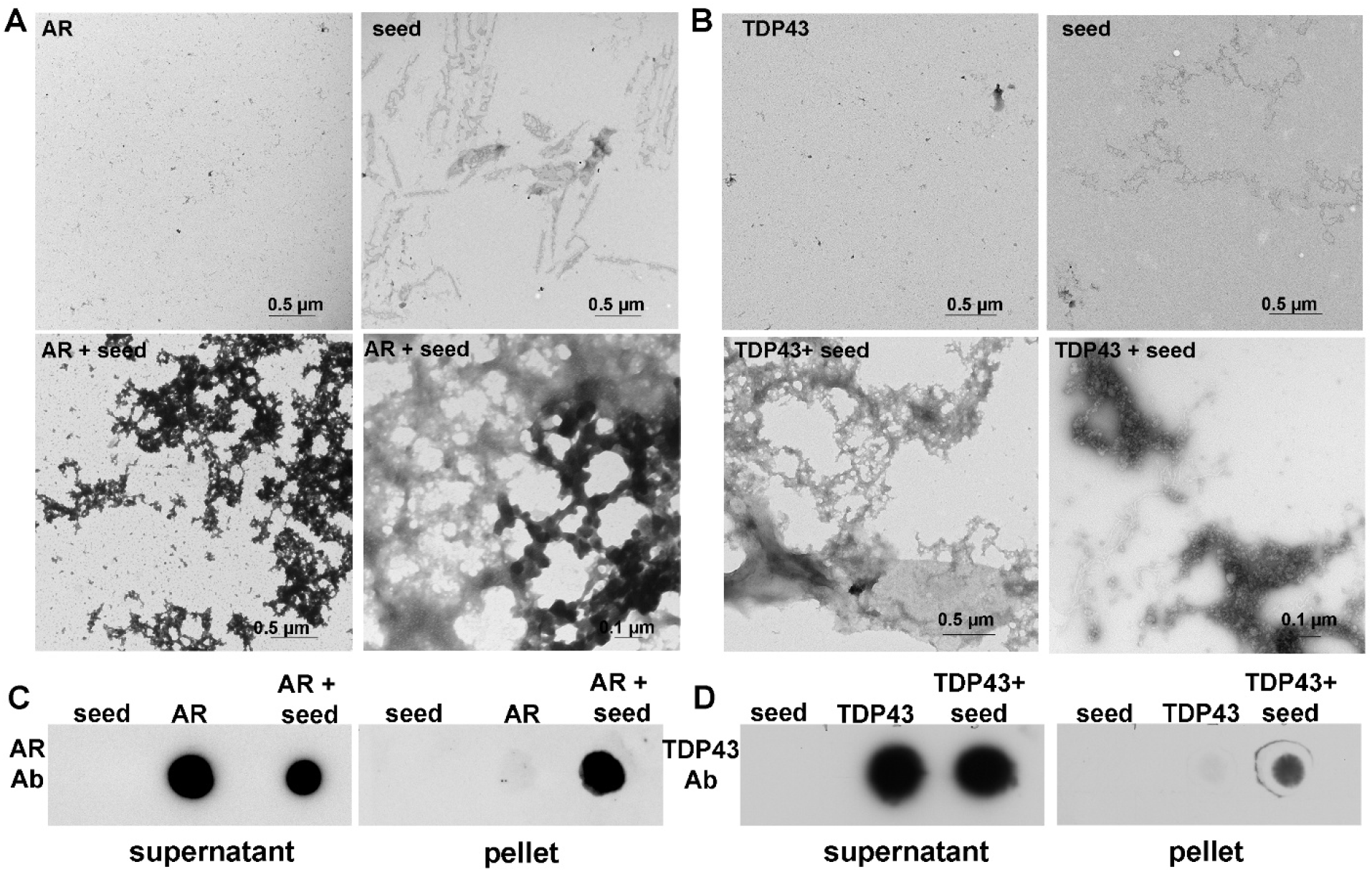
Perforatoria extracts template AR and TDP43 assembly. **A)** Negative stain transmission electron microscopy of seeding reactions of full-length rat AR alone (in the presence of SDS/DTT), perforatoria SDS/DTT supernatant seed alone, and AR templated by perforatoria seeds (AR + seed). Scale bar as indicated. n=3 rats. TEM of seeding reactions with other AR preparations is in Fig S3. **B)** Negative stain transmission electron microscopy of seeding reactions of full-length human TDP43 alone (in the presence of SDS/DTT), perforatoria SDS/DTT supernatant seed alone, and TDP43 templated by perforatoria seeds (TDP43 + seed). Scale bar as indicated. n=3 rats. **C)** AR and **D)** TDP43 seeding reactions were centrifuged to generate soluble (supernatant) and insoluble (pellet) fractions and examined for AR or TDP43 by dot blot analysis. n=2 rats.

Seeding reactions were also centrifuged to separate the resulting structures into soluble (supernatant) and insoluble (pellet) fractions and examined by dot blot using an anti-AR and anti- TDP43 antibodies. This experiment showed that while there were abundant AR or TDP43 monomer present in both the monomer alone and seeded reactions, only the reactions that received the perforatoria SDS/DTT seeds assembled into large AR positive or TDP43 positive structures that pelleted after centrifugation (Fig 11C,D). These results indicate the structures observed by TEM are constructed of AR or TDP43 and that the perforatoria extracts are needed to template their assembly.

While we anticipate the perforatoria seeds that served as templates were AR and TDP43 amyloids, they likely represent only a small percentage of the total perforatorium protein/amyloid content - as evidenced by the absence of signal of the seed alone reaction relative to that of recombinant proteins in dot blots- and therefore we cannot rule out the possibility of cross-seeding where perforatoria amyloids unrelated to AR and TDP43 served as templates for AR and TDP43 assembly. Nonetheless these experiments show that the cauda sperm perforatorium possesses amyloids that can propagate structure in naïve proteins.

## Discussion

Our studies presented here reveal that rat spermatozoa possess amyloid-rich germ granule-like condensates in the perforatorium, a cytoplasmic derived structure that is delivered to the oocyte during fertilization. While the perforatoria is unique to spermatozoa with a falciform (hooked) head (rodents, marsupials), the perinuclear theca, of which the perforatorium is a part, is a universal mammalian cytoskeletal structure found also in spatulate shaped sperm (human, domestic animals) suggesting germ granule-like condensates may also be in spermatozoa of other mammalian species. The rat perforatoria condensates contain highly conserved germ granule proteins and sncRNAs that play integral roles across species in genome protection, regulation of gene expression and germline specification. Although not proving function, the presence of both the executing protein and guide sncRNAs suggests the condensates can provide molecular information to alter the embryo’s RNA-regulatory state and possibly affect phenotype. We show also that perforatoria amyloids can transmit their structural conformations to naïve monomeric proteins. These results suggest a mechanism in rat spermatozoa allows information encoded in protein structure to be passed to naïve proteins in the embryo, i.e. protein-based inheritance. A central prediction of our model—that perforatorial condensates and their amyloid components are environmentally responsive sensors and transmitters of heritable information—was not directly tested here, as all experiments were performed in unperturbed normal SD rats. Using an established model for transgenerational epigenetic inheritance, studies are ongoing to determine if the rat sperm perforatoria condensates, and its amyloids, transmit heritable phenotypes to offspring of successive generations.

Although germ granule markers are also present in the amyloid-containing centrosomal region of the sperm tail, the highly ubiquitinated and insoluble properties of this region, abundance of lower molecular weight germ granule isoforms-suggestive of protein processing-, and lack of detectable RNA suggest this population represents remnants of the chromatoid body that may be associated with pericentrosomal aggreasomes for subsequent clearance through proteasomal degradation or autophagy. However, because centrosomal material can be delivered to the oocyte during fertilization, including in rodents, it is possible that conditions in which sperm quality control mechanisms are disrupted could result in aberrant proteins/amyloids being delivered to and affecting the developing embryo.

In most animals, germ granules are prominent during embryonic germ-cell specification or early germline development where they perform key functions in organizing post-transcriptional regulation of the germline program, serving as sites for RNA silencing, translational repression, RNA modification, and RNA turnover. As is typical for most biological condensates which assemble and disassemble based on cellular needs, the canonical cytoplasmic RNA–protein condensates (*C.elegans* processing (P bodies)/mammalian chromatoid body) do not survive spermatogenesis but instead are disassembled with most of the components likely eliminated through residual bodies. For this reason, germ granules are believed to be absent from spermatozoa of all animals and instead spermatozoa carry DNA methylation patterns, histone modifications, and distinct sncRNA cargo as the paternal information to the oocyte. Recent studies suggest, however, that the story is more complex than previously thought. In *C.elegans* new condensates are organized by BTB domain proteins (PEI-1 and PEI-2) that are trafficked to the developing spermatids where they recruit cytoplasmic argonaute WAGO3 and its associated 22G sRNAs (siRNAs) to function in mature spermatozoa as epigenetic carriers delivering new phenotypic and heritable information to the embryo ^20^. These sperm-specific PEI germ granules are distinct from the P-granules in the gonad not only based on function but also structure. Although the structural state of the PEI granules was not formally defined, the PEI granules exhibited resistance to 1,6-hexanediol, which dissolves liquid-like biomolecular condensates while WAGO3 showed slow exchange dynamics in FRAP indicating the PEI granules have more solid, possibly amyloid-like properties than the P body germ granule in the *C.elegans* gonad.

Our studies here suggest sperm-specific germ granule-like condensates are also present in rat spermatozoa. Indeed, perforatoria condensates are found in testicular spermatozoa, exhibit germ granule protein profiles distinct from that in the testicular germ cells, and are rich in amyloid indicating a more solid structural state than that of the chromatoid body which has the features of a viscoelastic RNP condensate^57^. Biological condensates and amyloids are now generally not viewed as separate states but as points along a continuum of protein self-assembly with increasing evidence that germ granules span a range of material states^58^. We speculate sperm- associated germ granules/condensates, as observed in *C.elegans* and here in rat, may exist in a more solid state to help maintain appropriate structural relationships/orientation during prolonged sperm storage prior to mating/fertilization, during the mechanical stress associated with the spermatozoon penetrating the oocyte and in early embryogenesis.

The rat perforatoria condensates also contained DNA/RNA binding proteins AR and TDP43 and their amyloid forms suggesting an additional level of regulation and complexity to condensate function. Beyond its established role as a nuclear hormone receptor and ligand activated transcription factor that mediates androgen signaling during spermatogenesis, the androgen receptor (AR) belongs to a family of polyglutamine (polyQ)-containing proteins that include Huntingtin. While pathological expansion of the AR polyQ tract promotes its aggregation and causes disease, aggregated AR has also been detected in the absence of polyQ expansion or overt pathology, raising the possibility that amyloid assembly contributes to normal AR function ^59–63^. These observations suggest that the functional state of AR may be coupled to its assembly into self-templating amyloid conformers, consistent with a prion-like mechanism of regulation. TDP-43 is a DNA/RNA-binding protein that regulates multiple aspects of RNA metabolism, including splicing, processing, stability, transport, and gene expression. Its low-complexity Q/N- rich prion-like domain enables reversible assembly into amyloid-like aggregates under conditions of cellular stress, resulting in altered protein function and cell state in a manner analogous to prion switching in yeast^64^. TDP-43 suppresses retrotransposition during early embryogenesis^65^ and promotes miRNA biogenesis as a component of the Drosha and Dicer complexes,^66^ highlighting essential roles in protecting genome integrity and posttranscriptional gene regulation. In the testis, TDP-43 functions as a transcriptional repressor in germ cells ^67^. While the full significance of AR and TDP43 in the sperm perforatoria condensates has yet to be determined, the presence of AR/AR amyloids in a germ granule-like structure may provide a mechanism for the germline to sense, respond to, and transmit to the embryo information of environmental hormones including endocrine disruptors with androgenic or antiandrogenic activity.

Rat perforatoria condensates, like germ granules, are enriched in proteins with sequence features (IDRs, low complexity, prion-like domains) that confer a high degree of environmental sensitivity enabling them to dynamically alter their structure and function in response to changes in their cellular environment. Indeed, the shape-shifting properties of amyloids is known including previous work from our lab where we showed an amyloid-rich mouse epididymal ECM adopted different amyloid conformations with different host defense functions depending on the bacterial strain^8,68,69^. Because of these inherent properties, we speculate phase-separating/amyloidogenic proteins may be the first responders to an environmental insult, encoding the biological information and their adaptation to the changed environment in their structures which then can be passed on to offspring via a prion-like propagation. If so, this would indicate a nongenomic protein- based mechanism of inheritance. In *C.elegans*, a functional amyloid was altered by the environment^20^ but other structural transitions are possible that could affect overall condensate structure/function. Germ granules and the rat perforatoria condensates also contain sncRNAs that function with their cognate proteins. Because of this relationship, changes in protein structure could also result in changes in sncRNA structure and function. In addition, whether sncRNAs can directly respond to the environment by changing structure is unknown, although a prion-like propagation of RNA/sncRNA has been proposed including as an epigenetic mechanism^70,71^. Indeed, it is possible that a combination of carriers (protein:sncRNA) can convey heritable information across generations.

The presence of germ granule condensates composed of environmentally sensitive proteins and sncRNAs with direct roles in modifying/protecting the genome may be a mechanism by which nature allows an organism to beta test new phenotypic changes before having them hard wired by changes in DNA sequence. The formation and consolidation of perforatoria condensates in the rat testis and epididymis, respectively, suggests there may be several opportunities to elicit change as alterations induced by exposures early in germ cell development could be enhanced/amplified or possibly corrected in the epididymis. During epididymal transit sperm sncRNA (miRNA, tsRNA) profiles change in part by their acquisition from epididymal extracellular vesicles^50,72,73^, and sperm-associated sncRNAs have been shown to transmit phenotypic information^74,75,76^. However, whether germ granule proteins are also delivered to epididymal spermatozoa is unknown. While much work is yet to be done, our studies suggest that mechanisms used by *C.elegans* to respond, and adapt to, a changed environment are conserved in mammals. Further studies are needed to establish perforatoria condensates/amyloids as bona fide paternal epigenetic carriers and their exact roles in transgenerational inheritance and the developmental origins of disease.

## METHODS

### Animals

All animal studies were conducted in accordance with the NIH Guidelines for the Care and Use of Experimental Animals using a protocol approved by the Texas Tech University Health Sciences Center Institutional Animal Care and Use Committee. Male Sprague-Dawley rats (∼90 days old) were purchased from Inotiv, Inc (West Layfayette, IN) (Harlan SD line) and maintained under a constant 12 hr light/12 dark cycle with food and water ad libitum. Rats were euthanized by CO_2_ asphyxiation followed by cardiac puncture. For all experiments, individual rats were the biological replicate except for RNA isolation where perforatoria were pooled from 5-6 rats.

### Isolation of testicular germ cells and epididymal spermatozoa

A mixed population of germ cells was isolated from the testis by removing the tunica vaginalis and testicular vein and mincing testis in a Petrie dish containing IMDM media to disperse the seminiferous tubules. The tubules were then transferred to a 50 ml centrifuge tube and allowed to settle for 15 min on ice. The top liquid was removed and the settled germ cells were treated with 10 ml of 1.5 mg/ml collagenase (Sigma #11088815001) in Dulbeccos PBS (dPBS) (Sigma #D5773) supplemented with 5 mM MgCl_2_ and 4 mM CaCl_2_ at 35^◦^C for 30 min while shaking (∼80 rpm). The dispersed germ cells were washed twice by resuspension in dPBS and centrifugation in a Beckman J6-MC rotor at 1500 rpm for 5 min at 4^◦^C. The germ cells were washed in 0.45% NaCl to remove red blood cells and then filtered through a 500 μm nylon mesh. The filtered cells were centrifuged at 1500 rpm for 5 min, 4^◦^C and the final pellet was resuspended in 10 ml cold dPBS. The samples were quantified by nanodrop, Laemmli buffer (LB) added to 1x final (62.5 mM Tris, pH 6.8, 6% glycerol, 2% SDS, 0.01’% bromophenol blue, 715 mM β-mercaptoethanol) and stored at -80^◦^C.

For isolation of testicular spermatozoa, tubules were minced in 20 mM phosphate buffer, pH 7.0 (buffer P, BP) containing 1x protease inhibitors (Roche cOmplete, Mini, EDTA-free Protease Inhibitor Cocktail, Sigma #11836170001) and 10 mM NaF and 80 mM Beta- Glycerophosphate (BGP)(Sigma #1183617000) (BP+ PI). The tubules were transferred to a 50 ml centrifuge tube and allowed to settle by gravity for 10 minutes on ice. The upper 5 ml of buffer containing testicular spermatozoa and germ cells (primarily elongating spermatids) was collected and centrifuged at 1500 rpm (466 x g), 4° C, 5 min in JS - 5.3 rotor, Beckman Avanti J-E centrifuge. Cells were resuspended gently in 3 ml of 0.5 x red blood cell lysis buffer (Invitrogen #00-4300-54) and then 3 ml BP + PI was immediately added and samples were centrifuged at 466 x g for 5 min. The cells were washed one time with 5 ml BP + PI and then spread on slides for immunofluorescence analysis.

To isolate epididymal spermatozoa, epididymides from one rat were pooled, cut into caput (subregions 1-3) or initial segment (subregion 1) and corpus-cauda (subregions 4-5 for all experiments except for the perforatoria condensate maturation studies, in which case only subregion 5 was used) and punctured using a 26G needle in BP+ PI. Caput and initial segment spermatozoa were allowed to disperse at room temperature for 10 min while cauda spermatozoa were dispersed by incubation at 37^◦^C for 30 min. Spermatozoa were centrifuged (1500 x g) and the supernatant was removed as luminal fluid and stored at -80◦C. Sperm pellets were washed twice in BP+PI, resuspended in 2 ml BP+PI (250 μl if from the initial segment) and concentration determined by counting with a hemacytometer.

### Isolation of sperm heads, tails and perforatoria

Spermatozoa in 2 mls buffer P + PI (∼ 1×10^8^ sperm/mL) were sonicated at 70% duty using a Heat Systems Ultrasonics W-375 microprobe, 45 seconds for cauda and 30 seconds for caput spermatozoa to detach sperm heads from tails. The heads and the tails were separated using sucrose density gradients following a procedure modified from Protopapas et al. ^28^ A 2.2 M stock solution of sucrose (Sigma Aldrich #S0389) in BP was prepared the previous day and gravity filtered. The gradients were prepared by layering 13 ml 2.2M sucrose on the bottom of an ultraclear centrifuge tube (Beckman # 344058) followed by 12 ml 2.05M and then 11 ml 1.85M sucrose containing the sample, which was made by adding 9 mls of 2.05M sucrose to the 2 ml sperm head/tail suspension. The gradients were spun in a Beckman Optima XE-90 ultracentrifuge using an SW32 rotor for 1hr 10min at 4^◦^ at 100,000 x g, using the slowest brake setting. The tails banded at the top of the 2.05 M sucrose, and heads formed a ring at the bottom of the tube. The tails were removed by puncturing the centrifuge tube with an 18G/ 1.5 inch needle/5 ml syringe, withdrawing the tail band up to 5 ml and bringing the sample up to 13 ml with BP. The tails were centrifuged at 10,000 x g for 20 minutes at 4^◦^C, the pellet resuspended in 1ml BP+PI, quantified by Nanodrop and stored at -80◦C. The remaining sucrose in the ultracentrifuge tube was removed by pipette down to ∼ 0.5 mls (as close to the ring of heads as possible without disturbing it). One ml of BP was added to the sperm head pellet and mixed gently to detach heads from the tube and transferred to a 15 ml centrifuge tube. An additional 1 ml BP was added to the ultracentrifuge tube and gently mixed to capture remaining heads. The ∼2.5 ml volume was brought to 5.5 ml with BP, 1.5 ml of the sperm head suspension was transferred to each of two 1.5 ml microfuge tubes and centrifuged at 20,300 x g for 20 min 4^◦^C using a Beckman JA18.1 rotor followed by a quick spin in a Shelton VS-15 microfuge at 15,000 rpm for 10 sec. The supernatant was discarded and the remaining head suspension was added to the existing pellets in the 2- 1.5 ml tubes and centrifugation steps repeated. The final two pellets were combined into 1 tube in a final volume of 500 μl BP + PI. The concentration of sperm heads was determined by counting with a hemacytometer.

For perforatorium isolation, sperm heads in 500 μl BP + PI (∼ 2×10^8^ sperm heads/ml) were transferred to low adherence microfuge tubes (USA Scientific #1415-2600) which was critical for optimal recovery of isolated perforatoria. The heads were sonicated for 4 bursts of 15 sec each at full duty, while the tube remained submerged in ice. After sonication, 250 μl of 80% sucrose (prepared in BP) was added, then the samples were brought up to 1 ml final volume with BP, which became the top layer (20% sucrose) for sucrose density gradients. The gradients were prepared by carefully layering 6 ml 80% sucrose on the bottom of an ultraclear centrifuge tube (Beckman # 344059) followed by 4 ml 30% sucrose and then 1 ml of the sucrose containing the sample. The gradients were spun in a Beckman Optima XE-90 ultracentrifuge using an SW41 rotor at 139,300 x g for 25 min at 4^◦^C, using the slowest brake setting. The perforatoria stopped at the top of the 80% sucrose and were removed by puncturing the centrifuge tube with an 18G/1.5 inch needle/5 ml syringe. Care was taken to not remove more than 1.7ml from the perforatorium gradient. If a higher volume is removed the sample will need to be diluted more in buffer P to allow perforatoria to pellet out of the sucrose. The samples were diluted to 9 ml in BP in a 15 ml tube and divided into two 1.5 mL low adherence tubes (∼1.5 ml/tube) and centrifuged at 25,500 x g for 20 min at 4^◦^C using a Beckman JA18.1 rotor, followed by a quick spin in a Shelton VS-15 at 15,000 RPM for 10 sec to repack the perforatoria. A repacking centrifugation step is needed since in low binding tubes the perforatoria pellet is not adherent and is easily disturbed when the tube is moved. One ml of the supernatant was quickly removed, the perforatoria repacked, 200 μl supernatant were removed, perforatoria repacked, and finally 100 μl more supernatant was removed. The remaining perforatoria in BP was added to the existing pellets and the process repeated until all perforatoria were pelleted out of the starting 9 mls. The final two perforatoria pellets were combined and concentration determined by Nanodrop.

### Indirect Immunofluorescence and ThS staining

Approximately 2-3 X 10^4^ intact epididymal spermatozoa, 4 X 10^4^ isolated heads and 0.18 mg tails were spread the day of isolation on Epredia Superfrost Plus slides (#6776214) in a 22 mm square and left to dry overnight. After rehydration in dPBS in a humid chamber, samples were fixed in 3.7% paraformaldehyde (Electron Microscopy Sciences, Hatfield, PA) for 30 minutes, permeabilized in 0.1% Triton X-100 (Thermo Scientific #28314) for 5 minutes, and rinsed thoroughly in dPBS-0.2% Tween-20 (Thermo Scientific #J20605-JP). Blocking buffer was made of 2% bovine serum albumin (Sigma Aldrich #A7511) in dPBS-0.2% Tween and applied for 30 minutes, except for the OC antibody for which 1% heat inactivated goat serum (Vector Labs, Newark, CA) was used. The primary antibody was diluted to 5 ug/ml in blocking buffer for all IgG purified antibodies, and 1:1000 for serum antibodies (see Table 1). The slides were incubated with antibodies for 1hr at room temperature, then overnight at 4^◦^C in a humid chamber. After washing in dPBS-0.2% Tween the secondary antibody was applied at 1:1000 in the blocking buffer and incubated for 1-2hrs in the dark at room temperature. Slides were washed in dPBS- 0.2%Tween and then incubated in 0.1mM DAPI (Molecular Probes, Invitrogen) for 10 mins at room temperature in the dark. Slides were washed in dPBS-0.2%Tween, rinsed in water, coverslips (Globe Scientific Inc, #1404) mounted with VectaShield (Vector Labs #H-100) and sealed with clear nail polish. All slides were imaged the day of, or the day after staining, and kept in the dark from the moment of sealing. For colocalization experiments, both primary and secondary antibodies were coincubated, respectively. For some experiments, tail samples were exposed to trypsin prior to antibody staining. Slides that were left to dry overnight were rehydrated in dPBS and then fixed in 4% paraformaldehyde for 15 mins at room temperature. They were washed in dPBS and incubated in 1 mg/mL Trypsin (Sigma #T-7418) for 3 min at 37^◦^ ^48^. Trypsin was removed by washing slides in dPBS and then put into blocking buffer followed by primary antibody as described above.

To examine condensate formation and maturation, testicular, initial segment (subregion 1) and cauda (subregion 5) epididymal spermatozoa were isolated as described above. Epididymal sperm were diluted 1:10 while testicular sperm were kept at the isolated concentration. Paraformaldehyde (Electron Microscopy Sciences) was added to 4% final and sperm suspensions allowed to fix for 15 min at room temperature. Spermatozoa were centrifuged at 2,000 x g for 5 minutes to remove fix followed by washing in BP + PI. Five μl were spread in a 20 mm square and left to air-dry for 10-15 mins (so that no visible droplets were left, but before the sample was completely dried). Samples were then permeabilized with 0.1 % Triton X-100 followed by blocking and antibody incubation as described above.

For the thioflavin S (Sigma #T1892) staining, slides of dried intact perforatoria or perforatoria extracts (described below) were submerged in 0.1% ThS (in water) in a Coplin jar in the dark for 2 hours. The slides were then washed three times in water, each wash lasting 10 minutes. After, three washes of 50% ethanol for 10 minutes each were done. The slides were then rinsed with water, mounted with VectaMount (Vector Laboratories #H-5501), coverslipped, and sealed with clear nail polish. Slides were imaged immediately using a Zeiss Axio Observer microscope equipped with excitation filter ∼380-400 nm and emission filter ∼425-450 nm to detect ThS yellow- green fluorescence.

### Imaging

Confocal images of germ granule marker localization within the perforatorium were captured as Z-stacks at 100x with a Zeiss LSM980 confocal microscope. Using the Zen 3.13 software’s 3D viewer, the sperm heads were angled to view underneath the perforatorium. To determine colocalization of proteins, immunofluorescence images taken with a Zeiss LSM980 confocal microscope were analyzed with Imaris 10.2.0 Coloc Module. Thresholds were set manually to try to remove dim nonspecific background signal. A colocalization channel showing the voxels that contain signal from both antibody channels was created using the thresholded images, and the percent colocalized voxels and Pearson’s correlation coefficient were averaged for all the images. Three replicates were done, and the final Pearson’s correlation coefficient reflects the average of the replicates.

### Proteinase K digestion

A 10 mg/ml stock solution of proteinase K (Promega #V302) was diluted to 1 ng/μl in 100 mM Tris buffer, pH 8.0. Two hundred pg PK was added to 1.5 x 10^6^ cauda heads or 25 ng recombinant full-length rat AR (prepared in-house) or 5 ng recombinant human TDP43 (ACRO Biosystems Group, #TD43 - H5145) in 30 ul 1X LB (∼6.7 pg/μl final) and incubated at room temperature for 0, 2 and 4 hrs. 4X LB was added to 1X final and samples immediately frozen and stored at -80^◦^C until analysis. For PK digestion of tails, 500 pg of PK was added to 50 μg of tails in 30 μl 1x LB buffer (16.7 pg/μl). After 1hr, 2hr, 3hr, 4hr, 4xLB was added to 1x final LB and stored at - 80 °C until analysis by immunoblot.

### Immunoblot analysis

Samples were heated for 5 min at 95^◦^C, separated on a 4 - 20% SDS-PAGE precast Tris-HCl Criterion gel (Bio-Rad Laboratories #3450032) or TGX Criterion gel (BioRad #5671093) in 1X Tris/glycine/SDS running buffer (Bio-Rad # 1610512) and transferred to PVDF membrane (Millipore, REF IPVH00010) using a BioRad Transblot system in 1X transfer buffer containing 25 mM Tris, 192 mM glycine, 10% methanol and 0.01% SDS. The membranes were blocked with 3% milk in TBS/0.2% Tween-20 (TBST) then incubated with 0.2-0.5 ug/mL primary antibodies (see Table 1) overnight in 3% milk/TBST with shaking in a cold room 4 °C. Blots were washed 3 times with TBST and then incubated for 2 hr at room temperature with HRP–conjugated secondary antibodies (Vector Labs) at 1:20,000 (v/v) dilution in 3% milk/TBST. After washing 6 times, 5 min each with TBST, the blots were exposed to West Pico PLUS (Thermo Scientific, #34580) or West Femto (Thermo Scientific, #34096) chemiluminescent substrate for 2-3 min before exposure to autoradiography film (Thomas Scientific, LLC.). For peptide blocking experiments, peptides were incubated with antibodies (5:1) in 3% blocking buffer at 4^◦^C overnight before incubating with blots.

### Protein Aggregation Disease pulldown

The protein aggregation disease (PAD) ligand kit was purchased from Microsens Biotechnologies, London, UK. Forty-eight-120 μg isolated perforatoria or 400 μg tails from cauda epididymal spermatozoa was pelleted and resuspended in 90% TC grade DMSO. Samples were vortexed and subsequently incubated for 90 min at room temperature. 100 μl BP+PI was added to each sample followed by 400 μl sterile water, 200 μl capture buffer, and 100 μl PAD reagent per the manufacturer’s protocol. After a 10 minute incubation at room temperature, 100 μl magnetic PAD beads were added and left for 90 minutes on a rotator at room temperature. Using a magnet to secure the beads, the amyloids bound to the beads were washed with PAD kit washing buffer #1 followed by two washes with PAD washing buffer #2. The beads were captured by magnet, all liquid was removed and the amyloids were eluted from the beads by incubation in 1x Laemmli buffer at 65^◦^C for 15 minutes. Samples were stored at -20^◦^C.

### Perforatorium and Differential Tail Extractions

Perforatoria were pelleted by centrifugation at 25,500 x g for 20 minutes and the pellet was resuspended in 1% SDS/2mM DTT/20mM Tris pH 9 ^77^ at a concentration of 2 mg/ml for Western blot or 0.5 mg/ml for seeding assays and extracted for 15 minutes on ice. The samples were centrifuged at 39,200 x g for 20 min at 4^◦^C to separate sup and pellet using a Beckman tabletop ultracentrifuge and TLA100 rotor. The supernatant was removed, aliquoted and stored at -80^◦^C. The pellet was resuspended in extraction buffer to the starting volume, aliquoted and stored at - 80^◦^C. For Western blot 20 μl each of supernatant and pellet were loaded/lane. For seeding 1 μl supernatant was used.

Protein content in isolated cauda tails in BP+PI was determined using the Thermo Scientific Nano Drop Lite Spectrophotometer. One mg sperm tails was centrifuged at 20,300 x g for 20 min at 4^◦^C. The tail pellets were resuspended in 400 μl extraction buffer 1 consisting of 150 mM NaCl, 50 mM Tris pH 7.4, 2 mM DTT, 10 mM NaF (Sigma #201154), 80 mM Beta- Glycerophosphate (BGP, Sigma #G9422) and protease inhibitors (Roche #11836170001). Samples were extracted on ice for 20 min, then centrifuged at 33,500 x g, for 20 min at 4^◦^C. The supernatant was collected and saved at -80^◦^C, labeled E1 extract. The pellets were resuspended in 400 μl extraction buffer 2, composed of buffer 1 plus 2% Triton X-100. Samples were extracted on ice for 20 min, centrifuged at 68,000 x g for 20 min at 4^◦^C. The supernatant was collected, saved at -80^◦^C, and labeled E2 extract. The pellets were resuspended in 400 μl of extraction buffer 3, composed of E1 buffer with 1% sarkosyl. The samples were extracted on ice for 20 min and centrifuged at 33,500 x g for 20 min at 4^◦^C. The supernatant was collected, saved at -80^◦^C, and labeled E3 extract. The pellets were resuspended in extraction buffer 4 which contained 7M Urea, 2% SDS, 2 mM DTT, 125 mM Tris pH 8, 10 mM NaF, and 80 mM BGP. The samples were incubated for 30 min at room temperature, followed by 10 bursts of sonication at 10% power level in a Sonicator Cell Disruptor W- 375 from Heat Systems- Ultrasonic, INC, and stored at -80^◦^C. Thirty μl of each extraction and 50 μg of starting nonextracted tails were examined by immunoblot.

### X-ray Diffraction

One hundred fifty μgs of pooled perforatoria isolated from two ∼90 day old SD rats were pelleted by centrifugation at 25,500 x g for 20 min in a Beckman JA18.1 rotor and resuspended in 25 μl 5 mM ammonium acetate. The next morning the sample was spun a second time and most of the supernatant was removed, leaving a slurry which was pulled up into a 0.7 mm quartz capillary tube. The sample was allowed to air dry in the presence of desiccant. X-ray diffraction data were acquired using a Rigaku Screen Machine Xray generator (Rigaku, The Woodlands, TX) utilizing CuKa radiation (1.5418 Å) with a focusing mirror (50kV, 0.6mA) and mercury CCDdetector. The distance from sample to detector was 75 mm. An in-house Python program was used to measure and calculate the average distance of the darkest points in each ring.

### Negative Stain Transmission Electron Microscopy (TEM)

Isolated perforatoria were spotted on to a formar-carbon coated grid (Ted Pella, Redding CA, # 01800N-F) for 5 minutes. After removing residual sample by wicking with filter paper, the grid was washed with water, stained with 2% uranyl acetate, and washed twice with water, each time using 5 ul for 1 minute, and wicking with filter paper to remove the liquid. The sample was then dried for 2 min and stored until analysis. Structures were imaged using a Hitachi 7640 microscope.

### RNA Isolation, Gel Electrophoresis, and RNA Staining

Immediately after completing perforatorium isolation, the perforatoria generated from 5-6 rats were pooled (∼ 250-450 μg) and total RNA isolated using the Invitrogen mirVana miRNA Isolation Kit (Invitrogen #AM1561) following the manufacturer’s protocol. The RNA was eluted in heated nuclease-free water (Invitrogen #46-2224), flash frozen in liquid N_2_ and stored at -80^◦^C. Three replicate experiments were done. Samples were sent to Admera Health (South Plainfield, NJ) for sncRNA sequencing and analysis. The RNA integrity number (RIN) for the three RNA preparations was 2.1, 2.2, and 2.0, typical for spermatozoa. A low RIN is a structural artifact of spermatozoa’s unique non-ribosomal RNA composition.

To examine RNA size distribution, an aliquot of RNA isolated from 230 μg of perforatorium from 3 rats, RNA isolated from heads from one rat, and RNA from heads without perforatorium from one rat, was mixed with an equal volume of 2X Novus TBE-Urea buffer, heated to 90^◦^C for 6 min and examined on a denaturing precast mini 15% PROTEAN TBE Urea gel (Bio-Rad #4566053). A microRNA Marker and 21-mer DNA oligonucleotide (BioLabs N2102S) were run as standards. The gel was stained for 12 min at room temperature with Sybr Gold nucleic acid gel stain (ThermoFisher #S11494) diluted 1:10,000 in TBE and imaged with a blue light transilluminator scanner.

Isolated perforatoria and sperm tails were stained with SytoRNA Select stain (Invitrogen #S32023). Perforatoria or tails were dried overnight on to Epredia Superfrost Plus slides (ThermoFisher #6776214), rehydrated the following day in dPBS and then fixed in cold 100% methanol at -20’C for 10 min. Slides were incubated with 0.5 μM SytoRNA Select in dPBS in the dark for 20 min at room temperature following the manufacturer’s instructions. Slides were washed in dPBS, mounted with VectaMount (Vector Labs), sealed with nail polish, and imaged immediately. After methanol fixation some tail slides were washed in dPBS and incubated in 1 mg/mL Trypsin (Sigma #T-7418) for 3 mins at 37^◦^C. Trypsin was removed by washing slides in dPBS followed by SytoRNA Select staining.

### Recombinant Androgen Receptor expression and purification

Constructs of full-length rat AR or rat AR lacking the ligand binding domain, both with an N- terminal His tag in the pET28 expression vector were prepared by Twist Bioscience (South San Francisco, CA). BL21- Gold (DE3) bacteria were transformed with plasmid DNA. Colonies were grown in 50 ml LB Broth Miller medium containing 100 μg/ml ampicillin overnight at 37° C with 250 rpm shaking for 16 hours. The overnight culture was used to start bacterial growth in 1 liter of Terrific Broth (TB) medium containing 100 μg/ml ampicillin and 0.4 % glycerol at 37° C until it reached an O.D 600 of 1.0 then the temperature was reduced to 22°C. IPTG (0.4 mg/mL) was added and the culture was allowed to grow for 20 to 22 hours with 250 rpm shaking in the incubator. The bacteria culture was centrifuged for 20 min, 4°C, 5000 rpm (6200 x g) in Beckman Coulter Avanti J – 26 XP, rotor JLA – 8.1 and pellets were frozen until use. Eight grams of bacteria pellets were resuspended in 40 ml of equilibration buffer containing 50 mM sodium phosphate buffer, 300 mM NaCl, 6 M guanidine-HCl, 20 mM imidazole, pH 7.4 and rocked in a cold room for 30 minutes followed by centrifugation for 30 min, 4°C, at 2190 x g. The supernatant was applied to a His60 Ni Superflow Resin (Clontech Laboratories, Inc. # 635660) column and incubated at room temperature for 30 minutes to 1 hour. The purification steps follow the manufacture’s manual protocol using gravity flow. The column was washed with 5 times column volumes of equilibration buffer and 10 times column volumes of wash buffer containing 50 mM sodium phosphate buffer, 300 mM NaCl, 6 M guanidine-HCl, 40 mM imidazole, pH 7.4. AR was eluted with elution buffer containing 50 mM sodium phosphate buffer, 300 mM NaCl, 6 M guanidine-HCl, 300 mM imidazole, pH 7.4. To remove imidazole and guanidine, the eluted protein was dialyzed 3 times with 2 liters of 1xPBS buffer pH 6.8 in a cold room. The final purified proteins were quantified by Nanodrop, aliquoted and stored at -80◦C. Protein identity was confirmed by immunoblotting (Fig S3).

HEK expi293F cells (ThermoFisher) were transfected with full-length rat AR (His-eGFP-AR in pTwist-CMV-BG-WPRE) using Epifectamine 293 transfection kit (Gibco/ThermoFisher #A14524). AR expression was induced using the Expi293 Expression system (ThermoFisher #A14635). Cells were harvested after 72 hrs and protein isolated by lysis of cells in 40 mM HEPES pH7.4, 300 mM NaCl, 1 mM TCEP, benzonase and protease inhibitors (Roche) using a parbomb to facilitate cell breakage. The lysate was centrifuged 12,000 x g for 20 min to separate soluble from insoluble fraction. The soluble supernatant was loaded onto a GFP nanobody column (a generous gift from Erik Yukl at New Mexico State University via R. Bryan Sutton, TTUHSC) and allowed to bind for 1 hr 4^◦^C. Beads were then washed by gravity flow with lysis buffer. AR was cleaved from its GFP tag while on the column using HRVC3 protease (a gift from R. Bryan Sutton lab) in an equimolar concentration to protein on the column incubating overnight at 4^◦^C. AR and protease were eluted from the column using lysis buffer. Sample was concentrated from ∼20 ml to 5 ml using an Amicon 10 kDa cutoff concentrator, loaded on to a Superdex 200 gel filtration column in 40 mM HEPES, 300 mM NaCl and AR protein collected.

### AR and TDP43 Seeding Reactions and Dot Blot Analysis

Five micrograms of full-length or N-terminal rat AR or three micrograms full-length human TDP- 43 (ACRO Biosystems #TD3-H5145, ACRO Biosystems, Newark, DE) were added to 1 μl rat perforatorium SDS/DTT extract supernatant (1% SDS, 2mM DTT in Tris buffer pH 9, 0.5 mg/ml) in a 96-well black flat bottom plate (Costar #3915, Corning Incorporated, ME) containing 20 µM ThT, 50 mM Tris pH 8, and 25 mg ± 3 mg Zirconia/Silica beads (BioSpec #11079101z, BioSpec Products, Inc., Bartlesville, OK) in a total volume of 100 µl. Monomer and seed alone reactions were done as controls with 1 ul SDS/DTT/Tris buffer pH 9 added to monomer in place of the perforatoria extract. Samples were done in duplicate. All reactions were incubated at room temperature for 3 hours with shaking at 282 rpm in a Cytation C10 microplate reader (Agilent Technology, DE). Five microliters of monomer alone, seed alone or seeding reaction were spotted on to formvar-carbon grids for TEM analysis while the remaining sample was used for dot blot analysis. Monomer, seed, and seeded reactions were spun at 72000 rpm using a TLA100 rotor in a tabletop ultracentrifuge, 4°C for 30 min to separate soluble (supernatant) from insoluble (pellet) material. The pellet was resuspended in 20 μl 50 mM Tris pH 8. Five µl of each supernatant and 20 μl of each pellet were manually spotted onto BioTrace nitrocellulose membrane (PALL Corporation, Mexico) using a dot blot apparatus (SRC-96 MINIFOLD, Schleicher & Schuell, Inc.) under vacuum but without the lid. The membranes were allowed to dry and then blocked with 3% milk in TBST (0.2 %Tween-20) for 1 hour at room temperature followed by incubation at 4^◦^C overnight with goat anti-human androgen receptor antibody (Abcam #19066, Abcam Inc., Waltham, MA) or rabbit anti-human TDP-43 antibody (Sigma #T1705). The membranes were washed in TBST 3 times, 5 minutes each then incubated with HRP-secondary antibodies (Vector Labs) for 2 hours at room temperature. The membranes were washed 6X in TBST, incubated with SuperSignal West Pico chemiluminescent substrate (ThermoScientific, Rockford, IL) followed by exposure to autoradiography film.

## Supporting information

Supplemental text for Figure 4

## Funding support

The John Templeton Foundation grant # 62184 (GAC).

## Acknowledgements

The authors gratefully acknowledge Mary Catherine Hastert for her assistance with the TEM analysis and Matthew Dominguez for assistance with AR expression.

## Notes

### Competing Interest Statement

The authors have declared no competing interest.

