## Supplemental text for Figure 4 for "Amyloid/RNA Condensates with Germ Granule Characteristics Are Present in Rat Epididymal Spermatozoa and Exhibit Prion-Like Propagation Activity"

Supplemental information:

Text

References

Table 1

Figs S1-S3

**Amyloid/RNA Condensates with Germ Granule Characteristics Are Present in Rat**

**Epididymal Spermatozoa and Exhibit Prion-Like Propagation Activity**

Kimberly Garza^1^, Uyen Tran^1^, R. Bryan Sutton^2^, Gail A. Cornwall^1*^

^1^Department of Cell Biology and Biochemistry and ^2^Department of Cell Physiology and Molecular Biophysics, Texas Tech University Health Sciences Center, Lubbock, TX 79430

Short title: Germ granule-like condensates in rat epididymal spermatozoa

*Corresponding author: Gail A. Cornwall, Ph.D., Department of Cell Biology and Biochemistry, Texas Tech University Health Sciences Center, 3601 4th Street, Lubbock, TX 79430..

Keywords: amyloid, epigenetic inheritance, germ granules, spermatozoa, rat, prion, RNA

Funding support: The John Templeton Foundation grant # 62184 (GAC).

**Supplemental text for Figure 4**

**DDX4/Vasa** is a DEAD box RNA helicase that is a core structural and functional component of germ granules. It acts as a key regulator of RNA within the granule with its primary function to control post-transcriptional gene expression. DDX4 associates with the piRNA pathway (PIWIs) to help suppress transposable elements and is essential for maintaining the germ cell lineage1. The full-length 79 kDa form of DDX4 along with isoforms of 55-60 kDa and ~ 45 kDa were detected in rat spermatozoa and isolated perforatoria. Similar DDX4 isoforms (79/80, 76/77, 55-60 and 37-45 kDa) were reported in the adult tamar testis, a marsupial, and predicted to also be in eutherian mammals.2 A DDX4 species at ~170 kDa may represent a stable dimer. All bands disappeared when the antibody was preincubated with its blocking peptide indicating antibody specificity (Fig S2).

**PIWIL1** functions primarily in post-transcriptional gene silencing by associating with piRNAs to form a piRNA-induced silencing complex (piRISC), which represses target transcripts, including transposon-derived RNAs, through mRNA degradation and translational repression. In addition to its role in RNA silencing, PIWIL1 contributes to chromatoid body assembly and is essential for maintaining germline integrity and specification.3 The canonical ~99 kDa PIWIL1 protein was detected in all samples. In addition to full-length protein, lower molecular weight PIWIL1-immunoreactive species, including bands at approximately 70–75 and 55 and 45 kDa, were also detected. The predominant species in the perforatoria were full-length and 70-75 kDa. A ~55 kDa PIWIL1 isoform lacking the MID and PIWI domains is predicted in the human genome (GeneCards). In addition, alternative PIWIL1 splice variants encoding C-terminally truncated proteins with altered PIWI domains, including predicted proteins of approximately 44 and 29 kDa, have been reported in bovine testis and ovary.4 All bands disappeared when the antibody was preincubated with its blocking peptide indicating antibody specificity (Fig S2).

**PIWIL4** plays a central role in transcriptional silencing of transposable elements. Loaded with piRNAs generated through the upstream PIWI pathway, PIWIL4 translocates to the nucleus, where it directs sequence-specific targeting of transposon transcripts and promotes de novo DNA methylation and repressive chromatin modifications at corresponding genomic loci 5 6. The full-length 96-kDa protein was the predominant species detected in all samples. Smaller isoforms that are C-terminal truncations of 56 and 52 kDa were detected in minor amounts and have been described in breast cancer cell lines.7 The even shorter forms of 23 and 13 kDa are indicated in Genecard and predicted as containing portions of the N-terminus. PIWIL4 immunoreactive forms of ~ 13 and 23 were detected only in the cauda sperm heads and isolated perforatoria. Unlike the tails, the perforatoria also contained several higher molecular weight species including those retained in the well suggestive of stable complexes (fuchsia bracket).

**TDRD7** is a member of the germline-enriched Tudor-domain-containing protein family. Tudor domains are conserved protein-interaction modules that can recognize methylated arginine or lysine residues. In male germ cells, TDRD proteins interact with arginine-methylated PIWI proteins, contributing to the organization and function of germline RNA-regulatory complexes8. TDRD7 regulates germ granule plasticity and the remodeling of RNA-protein complexes9. In addition to full-length TDRD7 (125 kDa) additional lower-molecular-weight TDRD7 species including ~60-, 50-, and 37-kDa forms, were detected in the rat spermatozoa. Similar isoforms (~70, 60, 50, 37, and 30 kDa) have previously been reported and attributed to alternative splicing in bivalves10 , while a ~60-kDa TDRD7 isoform has been described in human HEK293 cells 11 suggesting conservation of TDRD7 isoform diversity across species. The perforatoria also contained multiple higher molecular weight forms including those retained in the well (fuchsia bracket) suggestive of stable complexes.

**TDRD6** is a Tudor-domain-containing protein that plays a structural and organizational role in the chromatoid body12. TDRD6 also supports long 3′ UTR-triggered nonsense-mediated mRNA decay by facilitating the recruitment and assembly of mRNA-decay machinery within the chromatoid body.13 In addition, TDRD6 participates in early spliceosome maturation and splicing-related processes in primary spermatocytes14. In testicular germ cells and caput spermatozoa the canonical TDRD6 protein (~240 kDa) together with a ~200-kDa C-terminally truncated isoform previously described as enriched in the chromatoid body12 was observed. Cauda spermatozoa contained a higher-molecular-weight TDRD6 species (>250 kDa) whose detection varied between samples. Lower-molecular-weight immunoreactive bands of approximately 55, 37, and 17 kDa were also detected. Within the isolated perforatorium, the predominant TDRD6 species was the ~55-kDa protein, accompanied by the ~200-kDa C-terminally truncated and full-length TDRD6. All immunoreactive bands were markedly reduced or eliminated by preincubation of the antibody with its cognate blocking peptide, supporting their specificity for TDRD6 (Fig S2).

**BTBD18 and GMCL1 (BTBD13)** belong to the BTB/POZ domain-containing protein family. The BTB (Broad-complex, Tramtrack, and Bric-à-brac) domain is a conserved protein–protein interaction module that mediates homo- and heterodimerization and facilitates the recruitment of proteins into multiprotein complexes. In the testis, BTBD18 is a transcriptional regulator required for the expression of pachytene piRNA precursors at a subset of germline loci.15 The canonical ~78-kDa BTBD18 protein was detected in all samples. Testicular germ cells and spermatozoa also contained higher molecular weight (~110 kDa and >250 kDa), and smaller molecular weight (50 and 60 kDa) species. Similar proteins were observed using a second BTBD18 antibody and were eliminated following incubation of the antibody with the blocking peptide, supporting their specificity (Fig S2). Peptide competition also resulted in the appearance of new bands, including a 60–87-kDa smear, distinct from the canonical 78-kDa BTBD18 species, and a ~21-kDa band that may represent nonspecific interactions with the blocking peptide itself (Fig S2). In addition to canonical BTBD18, the perforatorium contained the ~60-kDa species and higher molecular weight assemblies including those retained in the well (fuchsia bracket).

**AGO2** is an miRNA/siRNA effector protein that functions as a core component of the RNA-induced silencing complex (RISC), mediating post-transcriptional gene regulation through miRNA-guided translational repression and mRNA decay.16,17 Unlike PIWI proteins, AGO2 is not germline-specific but associates with germ granule components and contributes to RNA silencing pathways in multiple cell types. The canonical ~97-kDa AGO2 protein was detected in all samples despite only labeling a discrete region at the base of the sperm head in immunofluorescence analysis suggesting inaccessible AGO2 epitopes. Smaller AGO2 species of approximately 55, 45, and 40 kDa were the predominant forms in spermatozoa, the insoluble sperm tail fraction, and the perforatorium. The ~55-kDa species may represent a halfAgo2-like product that lacks the C-terminal PIWI domain, analogous to a truncated AGO2 species previously described in human HEK293 cells18. In addition, noncanonical AGO2 forms have been reported in mouse oocytes, where a highly truncated AGO2 lacking essential functional domains and a predicted molecular weight of ~ 34 kDa suppresses miRNA activity during oocyte maturation 19. This smaller form was only detected in isolated cauda sperm heads and perforatoria.

**GMCL1 (BTBD13)** is a nuclear envelope-associated protein that may indirectly interface with germline RNA regulatory pathways20,21. Germ granules are frequently positioned near the nuclear envelope possibly to facilitate RNA surveillance 22,23. The canonical GMCL1 protein was found in testicular germ cells, spermatozoa, and in the insoluble fraction of isolated sperm tails. Although the predicted molecular weight of canonical GMCl1 is approximately 59–68 kDa, the protein exhibited anomalous migration on gradient SDS-PAGE, appearing closer to ~75 kDa. Additional lower-molecular-weight GMCL1 species were detected, with these forms enriched in the sperm tail fraction. In contrast, isolated sperm heads and perforatoria contained a predominant ~120-kDa GMCL1 species, together with an additional higher molecular weight immunoreactive form specifically observed in the perforatorium (fuchsia bracket).

**Dicer** is an RNase III enzyme that cleaves double-stranded RNA into small RNAs generating microRNAs (miRNAs) and small interfering RNAs (siRNAs) with a role in RNA silencing. 24 The canonical 219-kDa Dicer protein was in testicular germ cells and spermatozoa, together with a higher molecular weight species (>250 kDa) and several smaller isoforms (~145, 93, 60, 35, and 17 kDa). The 93-kDa isoform, which retains the RNase III catalytic domain and has been described as an alternatively spliced Dicer product in breast cancer cells25 was the predominant species detected in the perforatorium, along with full-length Dicer and an ~145-kDa isoform. In contrast, sperm tails were enriched for smaller Dicer species of approximately 35 and 17 kDa, which may represent proteolytic products, as well as several nonspecific bands that persisted following peptide competition of the antibody (Fig S2).
