## Supplemental Table 1 for "Amyloid/RNA Condensates with Germ Granule Characteristics Are Present in Rat Epididymal Spermatozoa and Exhibit Prion-Like Propagation Activity"

Table 1. Antibodies Used.

| **Antibody** | **Supplier/ Catalog Number** |
| --- | --- |
| Androgen Receptor | Abcam ab19066 |
| Androgen Receptor Peptide Block | Peptide 2.0 custom-made to Abcam antibody’s antigen |
| TDP43 | Sigma T1705 |
| A11 | Millipore AB9234 lot: NG1842996 |
| OC | Millipore AB2286 lot:3978216 |
| Speriolin | Generous gift from EM Eddy |
| FABP9 | Invitrogen PA5-122010 lot: YA3803133 |
| PIWIL1 | Novus NBP1-76489 |
| PIWIL1 Blocking Peptide | NBP1-76489PEP |
| PIWIL4 | Invitrogen PA5-116559 |
| AGO2 | Abcam ab32381 |
| DDX4 | Abcam AB13840 |
| DDX4 Blocking Peptide | Abcam AB13841 |
| BTBD18 | Invitrogen PA5-61933 |
| BTBD18 | Novus NBP2-14677 |
| BTBD18 Blocking Peptide | Novus NBP2-14677PEP |
| GMCL1 | Invitrogen PA5-58845 |
| GMCL1 Blocking Peptide | Antibodies Online ABIN6244258 |
| TDRD7 | Invitrogen PA5-70066 |
| TDRD6 | Novus NBP3-21396 |
| Dicer | Novus NBP2-88949 |
| Mono/Poly Ubiquitinylated Conjugates | Enzo ENZ-ABS840-0025 |
| Rabbit NRS | Jackson ImmunoResearch 011-000-120 |
| Rabbit IgG | Purified from Jackson ImmunoResearch NRS |
| Mouse IgG1 | Enzo ADI-SAB-600-0050 |
| Donkey anti Goat A-488 | Invitrogen A32814 |
| Donkey anti Rabbit A-594 | Invitrogen A21207 |
| Goat anti Mouse A-488 | Invitrogen A32723 |
| Donkey anti-Goat HRP | Invitrogen A16005 |
| Goat anti-Rabbit HRP | Vector Laboratories PI-1000-1 |
| Horse anti-Mouse HRP | Vector Laboratories PI-2000-1 |
