## Supplemental Figure 1 for "Amyloid/RNA Condensates with Germ Granule Characteristics Are Present in Rat Epididymal Spermatozoa and Exhibit Prion-Like Propagation Activity"

**Figures S1-3**

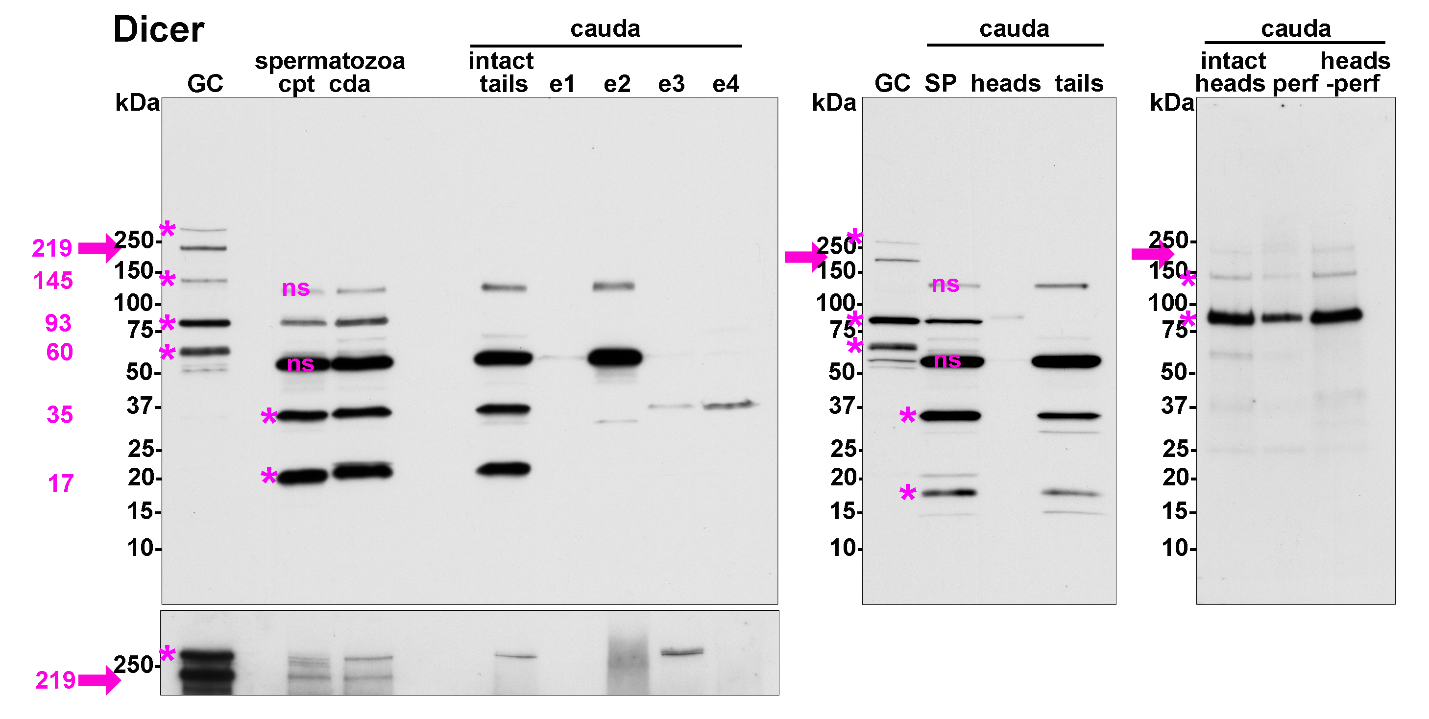

**Figure S1.** **Immunoblot analysis of Dicer in epididymal spermatozoa reveals protein profiles distinct from testicular germ cells and differences between sperm heads/perforatoria and tails.**

Fifty μg testicular germ cells (GC), 1 X 10^6^ intact caput (cpt) and cauda (cda) epididymal spermatozoa, 1.5 X 10^6^ isolated cauda sperm heads, 50 μg isolated sperm tails and 30 μg isolated perforatoria were loaded/lane. To examine solubility intact tails (50 μg) were sequentially extracted with E1, Tris buffer; E2, Triton X-100; E3, sarkosyl; E4, 7M urea. Canonical full-length protein and isoforms are indicated by fuchsia arrow and asterisks, respectively. Bottom panel, longer exposure of the blot revealing full-length Dicer. A blot incubated with peptide blocked anti-Dicer antibody is shown in Fig S2. ns, non-specific bands based on peptide blocking. n=2 rats.
