## Supplemental Figure 2 for "Amyloid/RNA Condensates with Germ Granule Characteristics Are Present in Rat Epididymal Spermatozoa and Exhibit Prion-Like Propagation Activity"

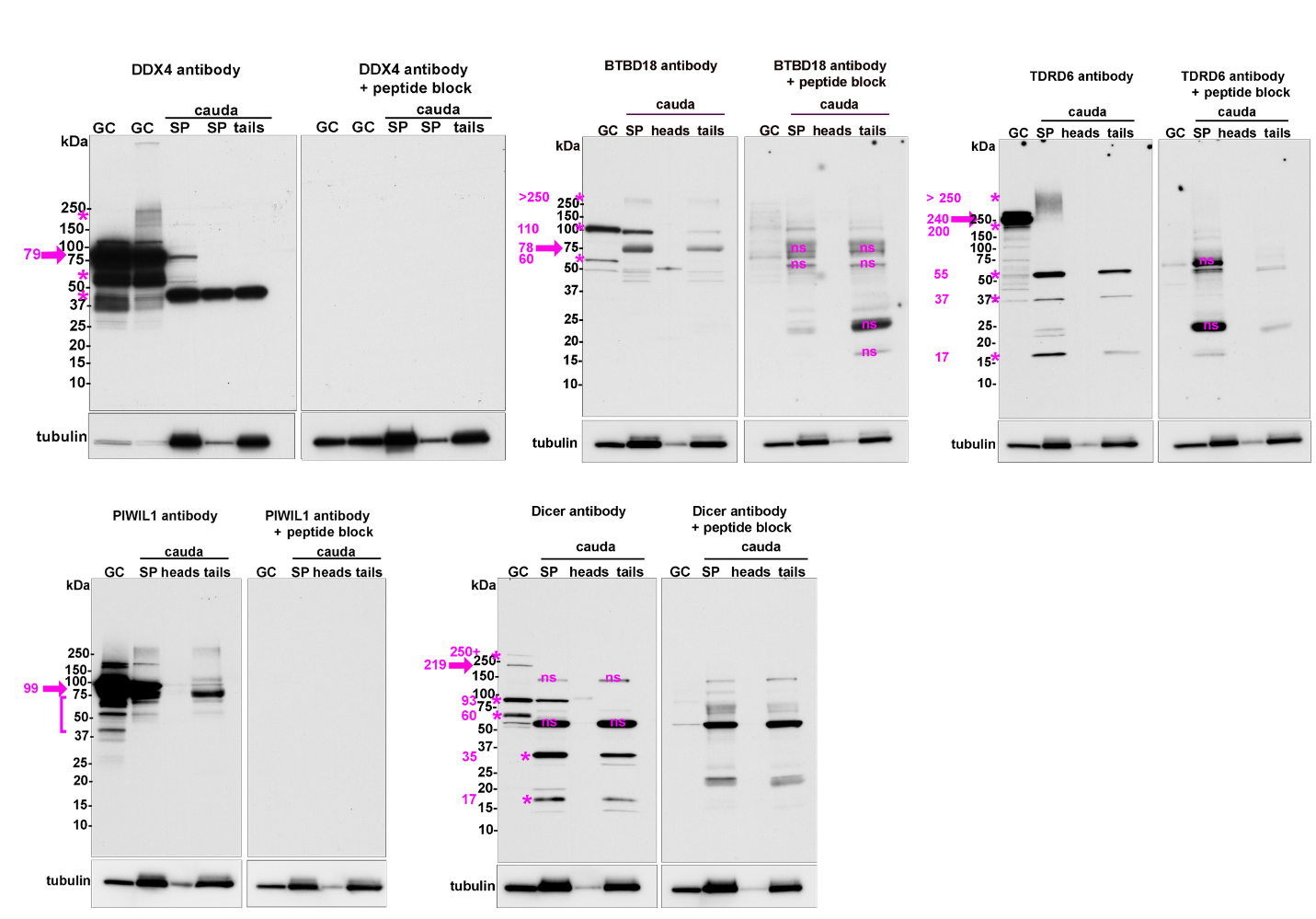

**Figure S2. Immunoblots with peptide blocked antibodies show antibody specificity.**

Fifty μg testicular germ cells (GC), 1 X 10^6^ intact caput (cpt) and cauda (cda) epididymal spermatozoa, 1.5 X 10^6^ isolated cauda sperm heads and 50 μg isolated sperm tails were loaded/lane. Blots were generated in parallel with one blot receiving the antibody and one receiving the peptide blocked antibody. Full-length proteins, fuchsia arrow; isoforms, fuchsia asterisks. ns, non-specific bands. n=2-3 rats.
