## Supplemental Figure 3 for "Amyloid/RNA Condensates with Germ Granule Characteristics Are Present in Rat Epididymal Spermatozoa and Exhibit Prion-Like Propagation Activity"

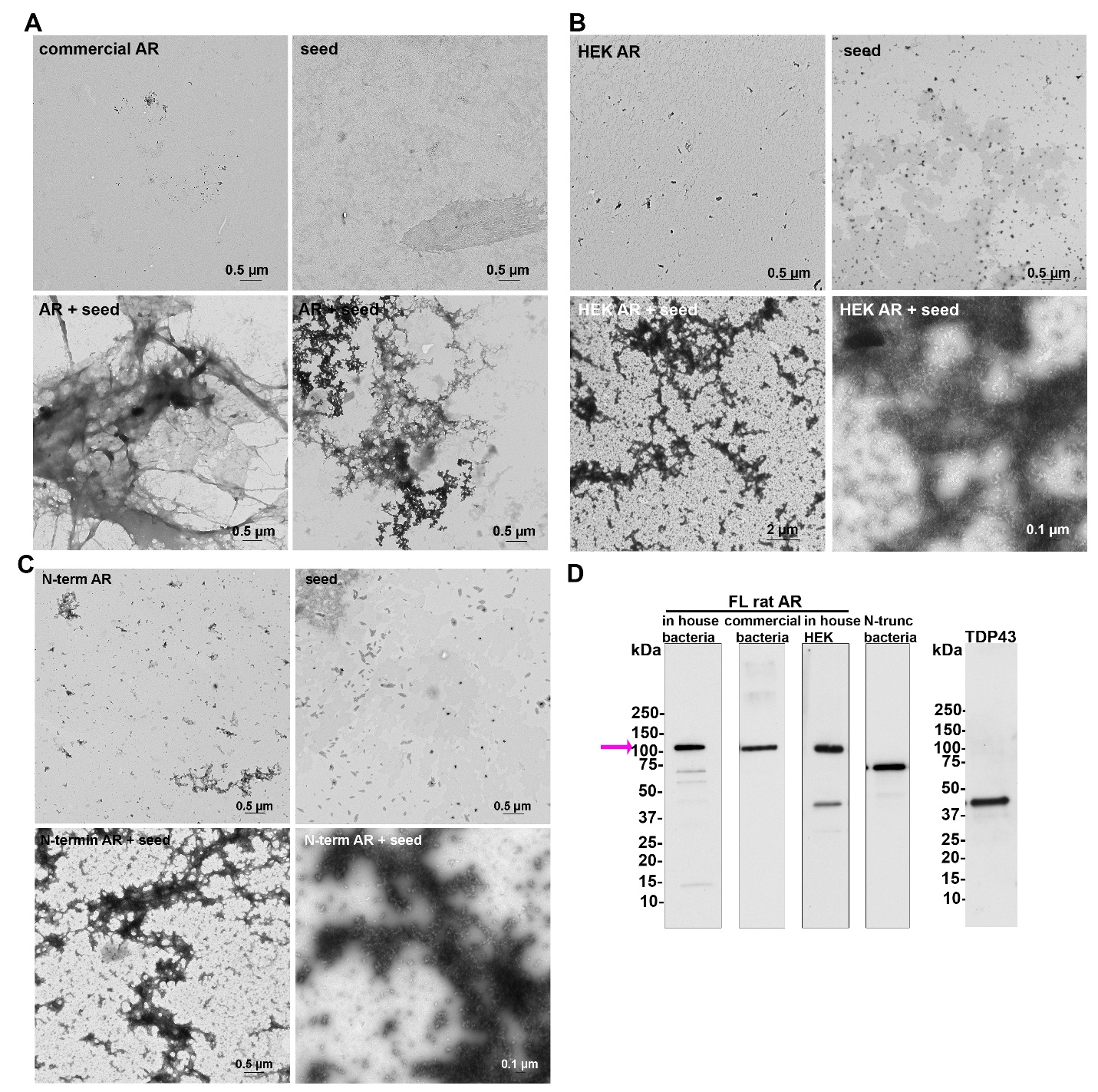

**Figure S3.**  **Perforatoria extracts template AR assembly.**

Negative stain transmission electron microscopy of seeding reactions of: **A)** commercially prepared full-length rat AR; **B)** AR expressed in HEK cels; and **C)** N-terminal AR lacking the ligand binding domain. For each, AR monomer alone (in the presence of SDS/DTT), perforatoria SDS/DTT supernatant seed alone, and AR templated by perforatoria seeds (AR + seed) are shown. Scale bar as indicated. n=2 rats. **D)** Immunoblot of recombinant AR and TDP43 proteins used for seeding assays. Full-length AR is indicated by fuchsia arrow.
